# Flexibility and constraint in preimplantation gene regulation in mouse

**DOI:** 10.1101/2020.03.13.990275

**Authors:** Colin C. Conine, Marina Krykbaeva, Lina Song, Robert C. Brewster, Nir Friedman, Oliver J. Rando

## Abstract

Although many features of embryonic development exhibit remarkable stability in the face of environmental perturbations, it is also clear that some aspects of early embryogenesis can be modulated by non-genetic influences during and after fertilization. Among potential perturbations experienced during reproduction, understanding the consequences of differing ex vivo fertilization methods at a molecular level is imperative for comprehending both the basic biology of early development and the potential consequences of assisted reproduction. Here, we set out to explore stable and flexible aspects of preimplantation gene expression using single-embryo RNA-sequencing of mouse embryos fertilized by natural mating, in vitro fertilization, or intracytoplasmic sperm injection, as well as oocytes parthenogenetically activated to develop (parthenotes). This dataset comprises a resource of over eight hundred individual embryos, which we use for three primary analyses. First, we characterize the effects of each fertilization method on early embryonic gene regulation, most notably finding decreased expression of trophectoderm markers at later stages of preimplantation development in ICSI embryos. Second, we find massive gene misregulation in parthenotes beyond the expected defects in imprinted gene expression, and show that many of these changes can be suppressed by sperm total RNA. Finally, we make use of the single-embryo resolution of our dataset to identify both stably-expressed genes and highly-variable genes in the early mouse embryo. Together, our data provide a detailed survey of the molecular consequences of different fertilization methods, establish parthenotes as a “tabula rasa” for understanding the role for sperm RNAs in preimplantation gene regulation, and identify subtypes of preimplantation embryos based on their expression of epivariable gene modules.

## INTRODUCTION

Development of multicellular organisms is a highly stereotyped process that begins with a single cell, the zygote, which proceeds through a set of choreographed stages. This process generates a multitude of differentiated cell types organized into a reproducible body plan. This process is extremely robust, occurring consistently in the face of extensive genetic variation across individuals of a given species. Moreover, proper development has proven to be robust to substantial environmental perturbations. For instance, expression of the developmental patterning protein Hunchback (Hb) is localized with remarkable precision (within one nuclear width) in *Drosophila* embryos at temperatures ranging from 9 to 29°C, despite extensive embryo-to-embryo variation in expression of its upstream regulator *bicoid* (Houchmandzadeh et al., 2002). These and many other features of development are heavily constrained by negative feedback control and other noise-reducing biological processes.

This robustness stands in contrast to the many sources of molecular variability that can drive cell-to-cell and organism-to-organism variation. At the single-cell level, statistical noise resulting from the small number of molecules present can (if not subject to stabilizing feedback) drive significant phenotypic variation across a population of cells (Raj and van Oudenaarden, 2008). Over longer timescales, so-called “epivariation” refers to stable maintenance of epigenetic information that in some cases can persist from one generation to the next. In mammals, a classic example of epivariation is the heritable variation in coat color observed in isogenic *A^vy^* animals, which is proposed to result from semi-stable replication of either wide or narrow cytosine methylation domains surrounding an IAP endogenous retrovirus (Morgan et al., 1999; Youngson and Whitelaw, 2008). Finally, transient environmental conditions can modulate organismal phenotypes that, in some cases, can persist long after the removal of the inciting insult. Among the most famous of such cases in mammals is the long-term reprogramming of offspring metabolism that occurs in response to poor nutrition experienced during gestation, a phenomenon widely studied as part of the “developmental origins of health and disease”, or DOHaD, field (Aiken and Ozanne, 2014; Simmons et al., 2001).

Here, we leverage low-input (in this case single-embryo) genomics to characterize flexibility and constraint in murine preimplantation gene regulation. In addition to characterizing gene expression variability across hundreds of nearly-isogenic embryos, we sought to characterize the responsiveness of early development to several early perturbations. Specifically, given that the details of fertilization method and embryo growth are generally believed to have at least some long-term impacts on development (Doherty et al., 2000; Mann et al., 2004; Rinaudo and Schultz, 2004; Rinaudo and Wang, 2012; Rivera et al., 2008; Vrooman and Bartolomei, 2017), we set out to identify persistent transcriptional changes driven by the different methods of fertilization used in assisted reproductive technologies (ART). Although assisted reproduction is generally considered safe (Halliday et al., 2019), it is clear that children conceived using assisted reproduction suffer from modestly increased risks for a wide range of disorders (Halliday et al., 2014). For example, intracytoplasmic sperm injection (ICSI) is associated with increased risks for congenital abnormalities, imprinting disorders, chromosomal abnormalities, and more subtle risks ranging from delayed neurological development to altered metabolism (Esteves et al., 2018). Given the length of the human lifespan and the recent (<50 years) development of ART, it is therefore of great biomedical importance to model assisted reproductive methods in other species to identify consequences for offspring physiology. Indeed, a number of groups have pioneered the use of rodent models to explore long-term sequelae of ART, demonstrating, for example, that even a single day of in vitro embryo culture can affect postnatal growth trajectory and metabolic parameters including glucose tolerance and fatty acid metabolism (Donjacour et al., 2014; Feuer et al., 2014; Giritharan et al., 2010; Giritharan et al., 2007; Riesche and Bartolomei, 2018; Vrooman and Bartolomei, 2017). In addition to these relatively well-characterized perturbations associated with in vitro embryo culture, assisted reproduction technologies also differ from natural mating in other ways (eg disruption of the zona pellucida by a piezo drill in ICSI).

Here, we gathered RNA-Seq data for hundreds of individual embryos, including embryos generated by natural mating (NM), in vitro fertilization (IVF), and intracytoplasmic sperm injection (ICSI). In addition, to develop a system for unbiased discovery of functions of sperm cargo in early gene regulation, we also characterized the transcriptome of hundreds of parthenotes – oocytes chemically activated to go through preimplantation development without any contribution from sperm – either left untreated or supplemented with sperm RNAs via microinjection. Our data provide insight into the effects of ART methods on gene expression in the embryo, identify widespread roles for sperm RNAs in control of early development, and reveal a surprisingly consistent landscape of gene expression flexibility in the preimplantation embryo.

## RESULTS

### Generation of matched sets of preimplantation embryos

To assay the early regulatory consequences resulting from assisted reproductive technologies, we generated four sets of embryos (**Figure 1A, Methods**). In all four cases, female mice of the FVB/NJ strain background were superovulated by staged injections of PSMG (Pregnant Mare Serum Gonadotropin) and hCG (human Chronionic Gonadotropin) 48 hours later. For natural mating (**NM**), a male was placed in a cage with a superovulated female immediately after hCG injection. Zygotes were collected the next day, stripped of cumulus cells, and subsequently cultured in vitro. For the three other conditions, eggs were collected from superovulated females and then treated as follows: 1) for in vitro fertilization (**IVF**), oocytes were incubated with motile cauda sperm allowing for fertilization in culture; 2) for intracytoplasmic sperm injection (**ICSI**), oocytes were fertilized by puncturing the oocyte’s zona pellucida using a piezo “drill”, followed by delivery of a sperm head into the ooplasm; 3) to characterize preimplantation development in the absence of any sperm contribution, we activated oocytes with strontium chloride to initiate parthenogenesis (**Parth**), treating them with cytochalasin B to prevent extrusion of the second polar body and thus maintain a diploid genomic complement.

**Figure 1.**
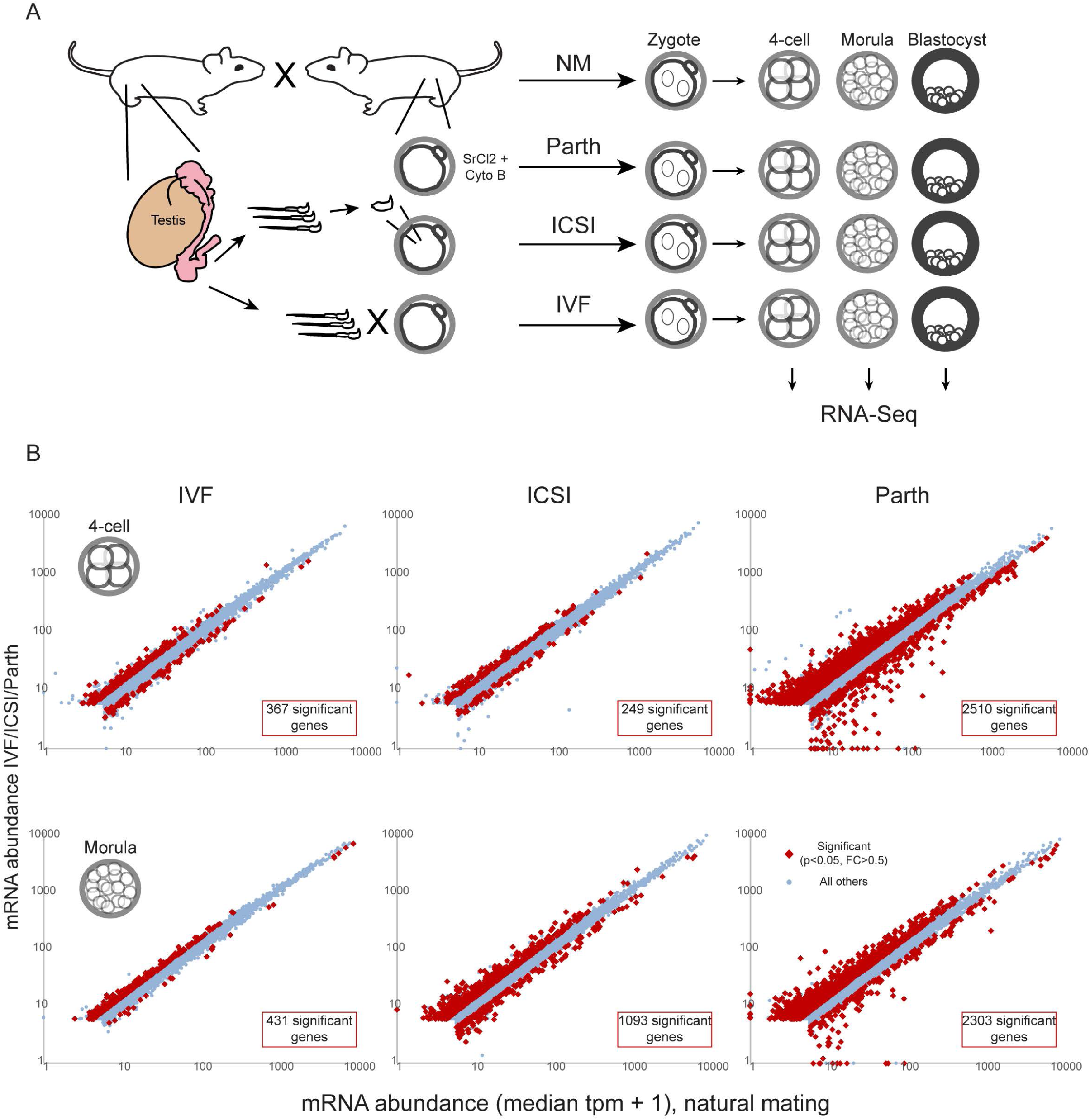
Schematic of the study and overview of the dataset. A) Cartoon of female superovulation and fertilization by 1) natural mating or 2) gamete collection with oocytes subsequently fertilized in vitro by IVF or ICSI, or activated parthenogenetically. Zygotes were then cultured to various stages and subject to single-embryo RNA-Seq. B) Scatterplots comparing mRNA abundance in embryos generated by natural mating (x axis) with embryos generated using IVF, ICSI, or parthenogenesis (y axis). For each scatterplot, the median mRNA abundance (tpm) was calculated across all embryos of a given type and values here are plotted as median + 1. Red dots show significantly (DESeq adjusted p value < 0.05, log_2_ FC > 0.5) misregulated genes for each comparison.

Importantly, several aspects of assisted reproduction that could potentially affect embryonic gene regulation are held constant across all four embryo sets. Most importantly, all four embryo sets were cultured in KSOM at 5% O_2_ – conditions which have previously been shown to minimize effects of ex vivo culture on preimplantation development (Doherty et al., 2000; Mann et al., 2004; Rinaudo and Schultz, 2004; Rivera et al., 2008) – thereby enabling comparison of all four embryo types under the same culture conditions and removing embryo culture vs. in vivo development as a variable. In addition, any lingering molecular effects of the superovulation process will be consistent across the entire set of embryos.

After fertilization (or activation), individual embryos were cultured to either the 4-cell stage (47 hours for parthenotes and ICSI embryos where the precise moment of fertilization is known, and 46 hours for NM and IVF where the time of fertilization is a 2-4 hour approximation) or the morula stage (75 hours post fertilization and just undergoing compaction) and collected for single-embryo RNA-Seq as previously described (Conine et al., 2018; Sharma et al., 2016). We also generated a smaller dataset for blastocyst-stage embryos, collected at roughly ∼96 hours but inspected for a typical ‘mid-blast’ appearance to minimize variability due to differing cell numbers. RNA-Seq libraries were generated using SMART-Seq2 (Ramskold et al., 2012), which captures full-length sequences across polyadenylated transcripts. Individual embryos were sequenced to a mean of ∼3.5 million reads per dataset (**Supplementary File 1**), and only embryos with at least 10,000 detected genes were used for further analysis. Complete data sets are provided in **Supplementary Files 2-3**, with an average of ∼60 embryos of each type for each stage.

As expected, these data robustly captured the major changes in gene expression between the four-cell and morula stages of development. Further supporting the quality of the dataset, male and female embryos were readily distinguished based on the anticorrelated expression of *Xist* and genes encoded on the Y chromosome (**Figure 1-figure supplement 1**). Comparing the various types of embryos generated here, we generally find that more dramatic perturbations to the fertilization process result in greater changes in gene expression: NM and IVF embryos were most similar overall (albeit not identical), ICSI embryos exhibited more extensive changes in gene expression, and parthenotes were clearly transcriptionally unique (**Figure 1B**). Below, we discuss the modest changes in expression between NM and both IVF and ICSI embryos, the more extensive gene expression differences specific to ICSI embryos, and finally, we use parthenogenesis as a system for understanding the role for sperm-delivered cargo in early development.

### Assisted reproduction affects expression of nuclear-encoded respiratory genes

We first focus on the relatively subtle differences between NM and IVF embryos (**Figure 1B**). Although NM and IVF embryos exhibited similar gene expression profiles overall, we identified 367 and 431 significantly-misregulated genes (adjusted p < 0.05, log_2_ fold change (FC) > 0.5) in IVF compared to the NM embryos at the 4-cell and morula stages, respectively. In terms of broad biological functions affected, GO enrichments in IVF-affected genes revealed significant downregulation of ribosomal protein genes and other genes involved in ribosome biogenesis, along with upregulation of genes involved in chromatin modification pathways, in IVF embryos at both 4-cell and morula stages. To determine whether this gene expression profile results from specific features of the IVF protocol, or simply the difference between fertilization in vivo and ex vivo, we compared gene expression in NM embryos to both IVF and ICSI embryos. Notably, IVF and ICSI effects on gene expression were positively correlated both for 4-cell and morula-stage embryos (**Figures 2A-B**), arguing that these transcriptional changes reflect the difference between fertilization (and its immediate aftermath) occurring in the female reproductive tract, vs. fertilization in culture.

**Figure 2.**
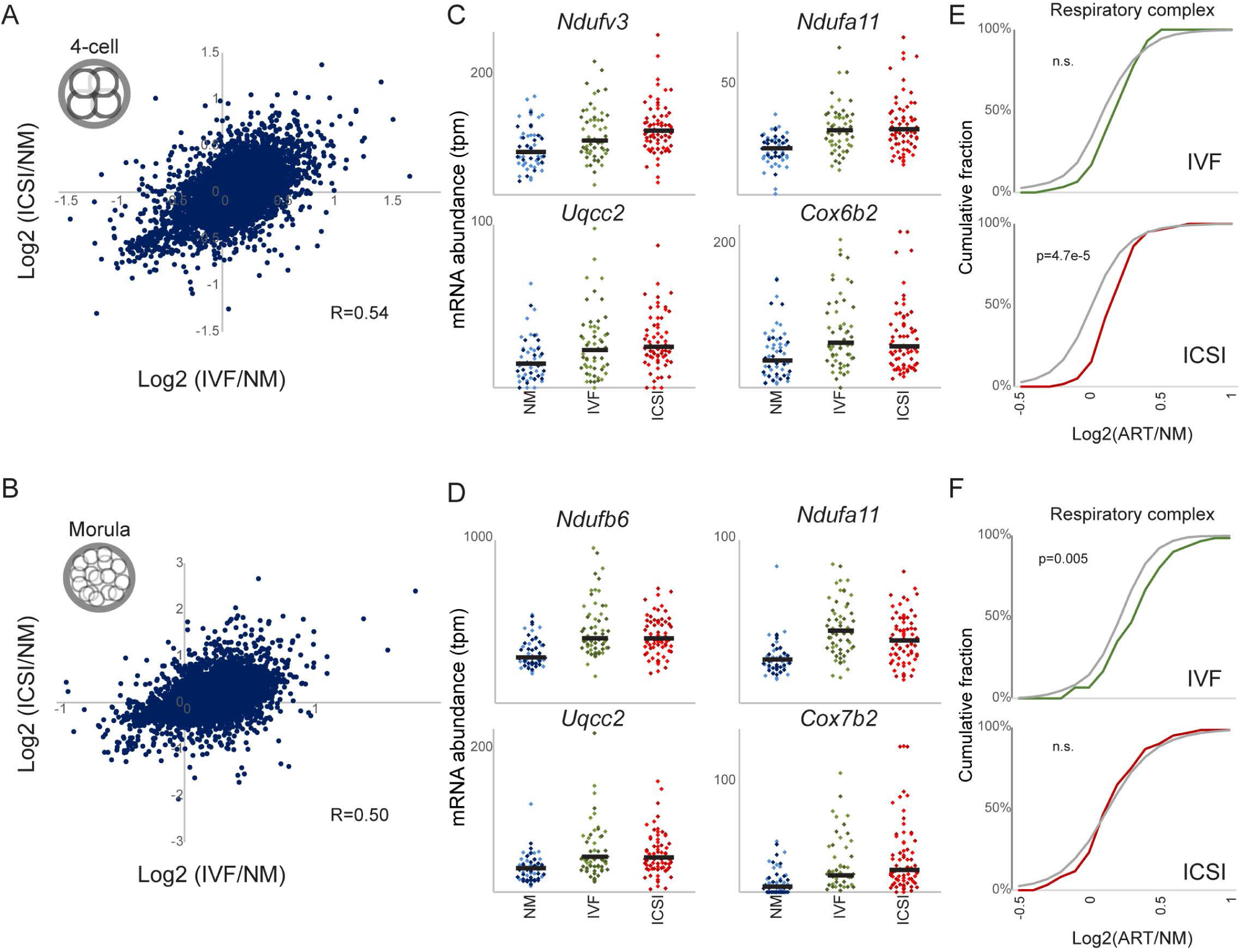
Overexpression of nuclear-encoded mitochondrial genes in assisted reproduction embryos. A-B) Effects of IVF and ICSI relative to NM are well-correlated. X axis shows the effect of IVF relative to NM as log_2_FC (IVF+1/NM+1) with y axis showing the equivalent effects of ICSI. Effects of assisted reproduction were observed both in 4-cell-stage (A) and morula-stage (B) embryos. C-D) Dot plots for key respiratory chain genes in individual NM, IVF, and ICSI embryos, as indicated, for four-cell-stage (C) and morula-stage (D) embryos. Here, each dot represents the mRNA abundance for one individual embryo. E-F) Cumulative distribution plots for changes in expression of genes annotated with the GO category “respiratory complex”. In each case, colored line shows the cumulative distribution of the effects of assisted reproduction (e.g. log_2_(IVF/NM) or log_2_(ICSI/NM)) across all annotated respiratory complex genes, with the gray line showing distribution for all other genes. See also Figure 2-figure supplement 1.

Focusing on shared biological functions affected by both ICSI and IVF, we confirmed that both IVF and ICSI embryos exhibited downregulation of ribosomal protein genes, although this was confined to morula-stage embryos for the ICSI cohort (**Figure 2-figure supplement 1**). We also noted a subtle enrichment of mitochondria-related pathways in a subset of the IVF and ICSI datasets, which motivated further exploration given prior reports documenting overexpression of genes involved in mitochondrial function (*Ndufa3*, *Cox6b2*, etc.) in ICSI blastocysts (Giritharan et al., 2010). Indeed, we find that multiple nuclear-encoded genes encoding respiratory chain components (*Cox7b2*, *Ndufa1*, *Ndufa11*, *Tomm6*, *Atp1b1*, *Ndufb4*, *Mrps9*, *Uqcr11*, *Ndufa2*, *Mrpl57*, etc.) were modestly overexpressed in both IVF and ICSI embryos relative to NM embryos (**Figures 2C-F**).

Together, these data demonstrate that the precise method of fertilization can have persistent effects on early development. Beyond the shared effects of IVF and ICSI on expression of genes involved in ribosome biogenesis and mitochondrial function, many other genes were consistently affected in these two cohorts (**Figures 2A-B**), although we found little compelling evidence for other clear biological functions affected in this shared gene set.

### Intracytoplasmic sperm injection results in altered expression of trophectoderm markers

Compared to NM and IVF, ICSI is a substantially more artificial method of fertilization. Most notably, the ICSI procedure involves physically disrupting the zona pellucida with a piezo “drill,” in contrast to the sperm acrosome reaction which serves as the natural mechanism for penetrating the zona. In addition, while the entire contents of the sperm are typically delivered to the oocyte in NM and IVF, when performing ICSI in mice only the sperm head is injected into the oocyte. Moreover, delivery of the sperm head is associated with a transient physical disruption to the oolemma, potentially resulting in leakage of oocyte contents.

We set out to identify abnormalities specific to the ICSI process by comparing ICSI and IVF embryos. To confirm and extend our initial findings (below), we also generated an additional replicate set of IVF and ICSI 4-cell- and morula-stage embryos (**Supplemental Files 4-5**, **Figure 3-figure supplements 1**-**2**). In this iteration, we generated two groups of ICSI embryos for each stage – a control group was microinjected with GFP mRNA as a marker for successful microinjection, while the experimental group was also injected with piRNAs (28-32 nt RNAs gel-purified from murine testis RNA). This experiment was motivated by our prior finding that piRNAs in mature sperm are relatively enriched in the sperm tail and depleted from sperm heads (Sharma et al., 2018), and are therefore lacking in typical murine ICSI protocols which rely on sperm head injection. Overall, comparing our first and second cohorts of embryos, the specific effects of ICSI on preimplantation gene regulation were moderately reproducible at the 4-cell stage (R=0.33) and highly reproducible at the morula stage (R=0.74) (**Figure 3-figure supplement 2**). Importantly, microinjection of testicular piRNAs had no effect on gene expression in either 4-cell- or morula-stage embryos (**Figure 3-figure supplement 1**), arguing against a major biological function of piRNAs in the zygote (at least as reflected in mRNA-Seq data) while simultaneously providing further confidence in the reproducibility of our datasets.

**Figure 3.**
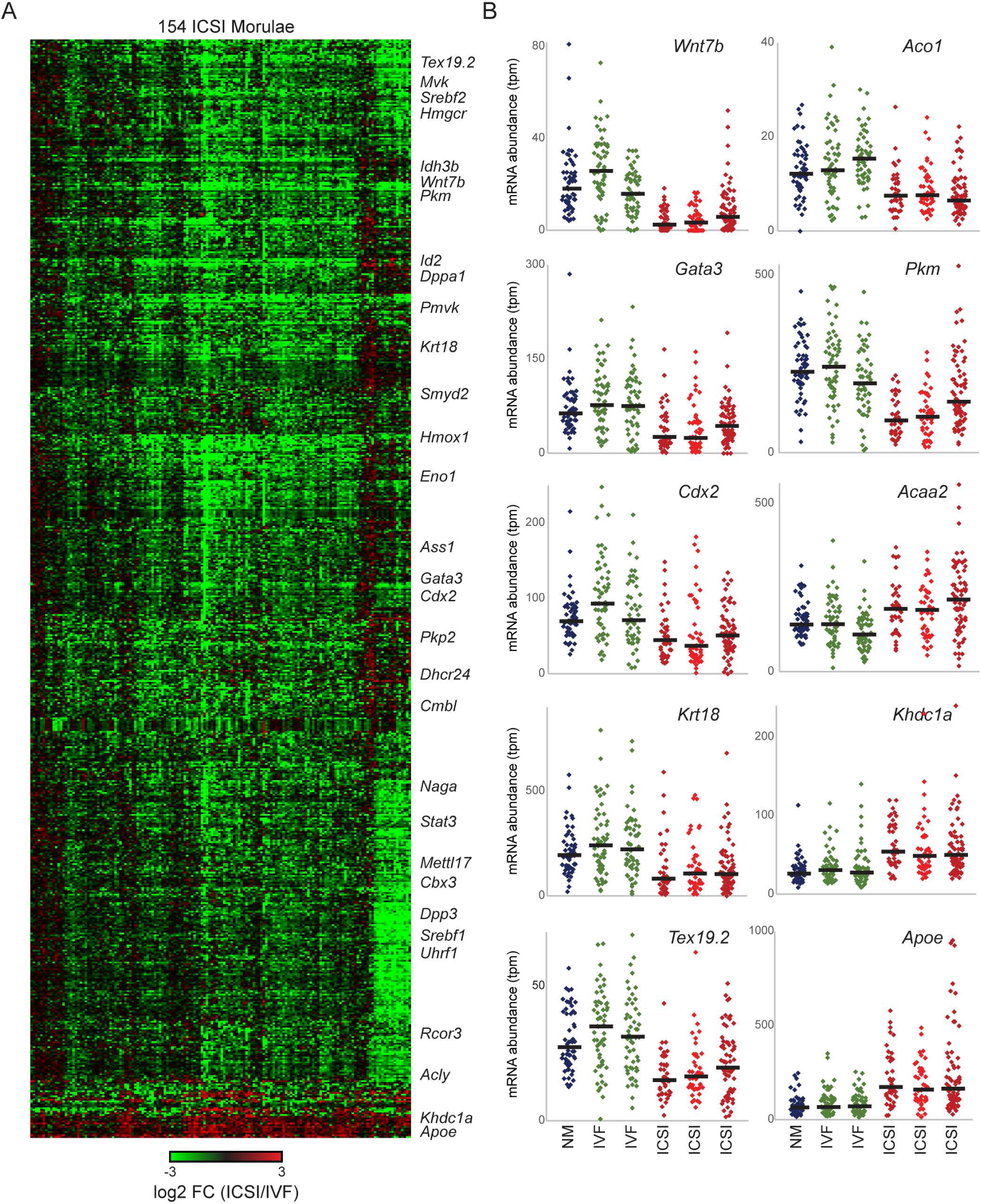
Widespread misregulation of trophectoderm markers in ICSI embryos. A) Heatmap shows significantly (adjusted p < 0.05) misregulated genes in all individual morula-stage ICSI embryos with each column showing the expression level for an individual ICSI embryo relative to the median expression level for the IVF embryo data set for each gene. B) Dot plots show individual embryo data for key trophectoderm markers and various metabolic enzymes. Data shown here include the original set of three embryo types (NM, IVF, ICSI) along with data from a second replicate series with one IVF cohort and two ICSI cohorts (injection of control (GFP) RNA or purified piRNAs into the zygote). See also Figure 3-figure supplements 1-3.

As our morula-stage data were more robust and biologically coherent, we focus on this stage for further analysis of ICSI effects on gene regulation. The vast majority of genes misregulated as a result of IVF were repressed in ICSI relative to IVF embryos (**Figure 3A**). Genes downregulated in ICSI embryos included a number of metabolic factors (*Aco1*, *Pkm*, *Acaa2*), along with well-known markers of the trophectoderm lineage such as *Cdx2*, *Gata3*, and *Krt18*. Not only were these genes consistently downregulated in three distinct morula-stage ICSI cohorts (**Figure 3B**), but we further confirmed downregulation of trophectoderm markers in a small blastocyst-stage dataset (**Figure 3-figure supplement 3**). Thus, although it is plausible that TE downregulation simply reflects differences in developmental progress between ICSI and IVF/NM embryos, the downregulation of these genes at two distinct developmental stages argues against this interpretation.

Importantly, as trophectoderm cells ultimately develop into the placenta and other extraembryonic tissues, our data may provide clues as to the origins of metabolic sequelae – most notably, deficits in glucose control – which have previously been documented in ICSI offspring (Rinaudo and Wang, 2012). Our findings suggest that altered trophectoderm cell number or function could potentially affect placental development, and consequently, fetal nourishment after implantation.

### Parthenogenesis results in widespread regulatory aberrations

We next turned to our parthenote dataset to characterize gene expression differences between fertilized embryos and embryos developing in the absence of any sperm contributions. Although it is well-understood that parthenotes exhibit widespread regulatory aberrations such as failure to express maternally-imprinted genes (Inoue et al., 2017a; McGrath and Solter, 1984; Surani and Barton, 1983), parthenotes do efficiently progress to the blastocyst stage, allowing their use as an experimental tool to interrogate the functions of sperm-delivered molecules in the early embryo. Consistent with prior genome-wide analyses of imprinted gene misregulation in parthenotes (Inoue et al., 2017a), we recapitulated the widespread loss of expression of maternally-silenced genes in the absence of a functional paternal genome (**Figure 4A**). These maternally-silenced genes include the X inactivation factor *Xist* (**Figure 4B**) (Inoue et al., 2017b), resulting in the expected loss of X chromosome dosage compensation at later stages of preimplantation development (**Figures 4C-D**). Overall, the majority of genes exhibiting complete loss of expression in the absence of a paternal genome were previously known (Inoue et al., 2017a), although we did identify a small number of apparently novel maternally-silenced genes, such as the lncRNA *Lincenc*.

**Figure 4.**
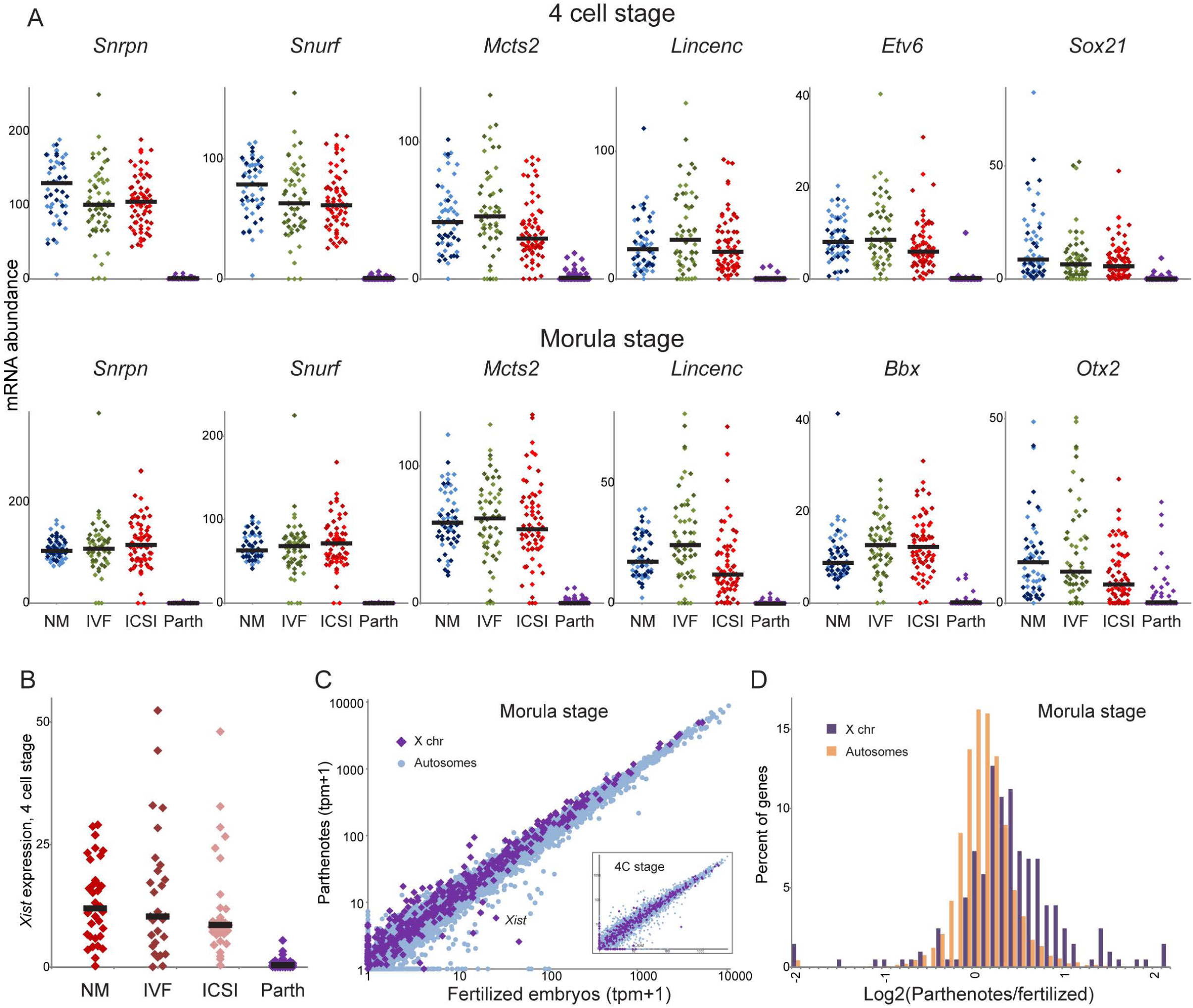
Misregulation of imprinted genes in parthenotes. A) Dot plots, as in Figures 2C-D and **3B**, showing data for known and novel maternally-silenced genes in all four classes of embryos in this study. As expected, maternally-silenced genes are not expressed in the absence of a paternal genome. For all fertilized embryos, data for male and female embryos are shown in different shades of blue/green/red, but none of these genes exhibit sexually-dimorphic expression. B) As in panel (A), showing loss of imprinted *Xist* expression in parthenotes. C) Scatterplot showing all genes (light blue) and X chromosome-encoded genes (purple), comparing fertilized embryos (x axis) with parthenotes (y axis). Main panel shows data for morula-stage embryos, inset shows 4-cell-stage data. D) Histogram of gene expression changes in parthenotes (log_2_ parthenotes/fertilized embryos) for X-linked and autosomal genes, as indicated.

Beyond the widespread deficits in imprinted gene expression, thousands of additional genes were misregulated in parthenotes (**Figure 1B**). Although many of these genes presumably exhibit altered expression as a secondary consequence of defective imprinting (eg genes affected by the absence of *Mcts2* expression, etc.), these genes also potentially include targets of sperm-delivered regulatory factors. For example, we have previously shown that an abundant cleavage product of tRNA-Gly-GCC (“tRF-GG”) present in sperm drives repression, in both ES cells and early embryos, of a group of genes regulated by nearby copies of the MERVL endogenous retroelement (Sharma et al., 2016). Consequently, these MERVL-regulated genes should be overexpressed in embryos deprived of sperm-delivered tRF-GG. Consistent with this prediction, previously-identified targets of tRF-GG were broadly overexpressed in parthenotes (**Figure 4-figure supplement 1**), providing further evidence supporting the proposed role for sperm-delivered RNA in early gene control.

Encouraged by the potential for parthenotes to recapitulate at least some early regulatory functions of sperm RNAs, we set out to empirically identify genes regulated by sperm RNAs in the early embryo. We purified sperm total RNA for microinjection into parthenotes, and compared matched sets of control-injected and sperm RNA-injected parthenotes by single-embryo RNA-Seq at the 4-cell stage (**Figure 5A, Supplemental File 6**). We first focused on two groups of genes we previously identified as targets of sperm-delivered small RNAs. One group consists of the above-mentioned targets of tRF-GG (Sharma et al., 2016). The second group comprises RNA and chromatin-related genes repressed by microRNAs – present in genomic clusters and thus dubbed “clustermiRs” for brevity – that appear to be absent from caput epididymal sperm but present in more mature cauda epididymal sperm (Conine et al., 2018, 2019). Consistent with predictions from these prior studies, we found that sperm total RNA injections into parthenotes led to repression of both tRF-GG- and clustermiR-regulated genes (**Figures 5B-C**). Beyond these predicted sperm-responsive genesets (together comprising fewer than 100 target genes), we documented more extensive effects of sperm total RNA on gene regulation in parthenotes. Indeed, we found a remarkable correlation between the effects of fertilization on mRNA abundance and the effects of sperm total RNA on parthenote gene regulation (**Figure 5D**), suggesting that sperm RNA can account for at least ∼10% of the gene expression changes in parthenotes. This finding thus identifies hundreds of genes regulated by the sperm RNA payload – presumably including both direct targets and transcripts affected as secondary consequences downstream of the direct targets.

**Figure 5.**
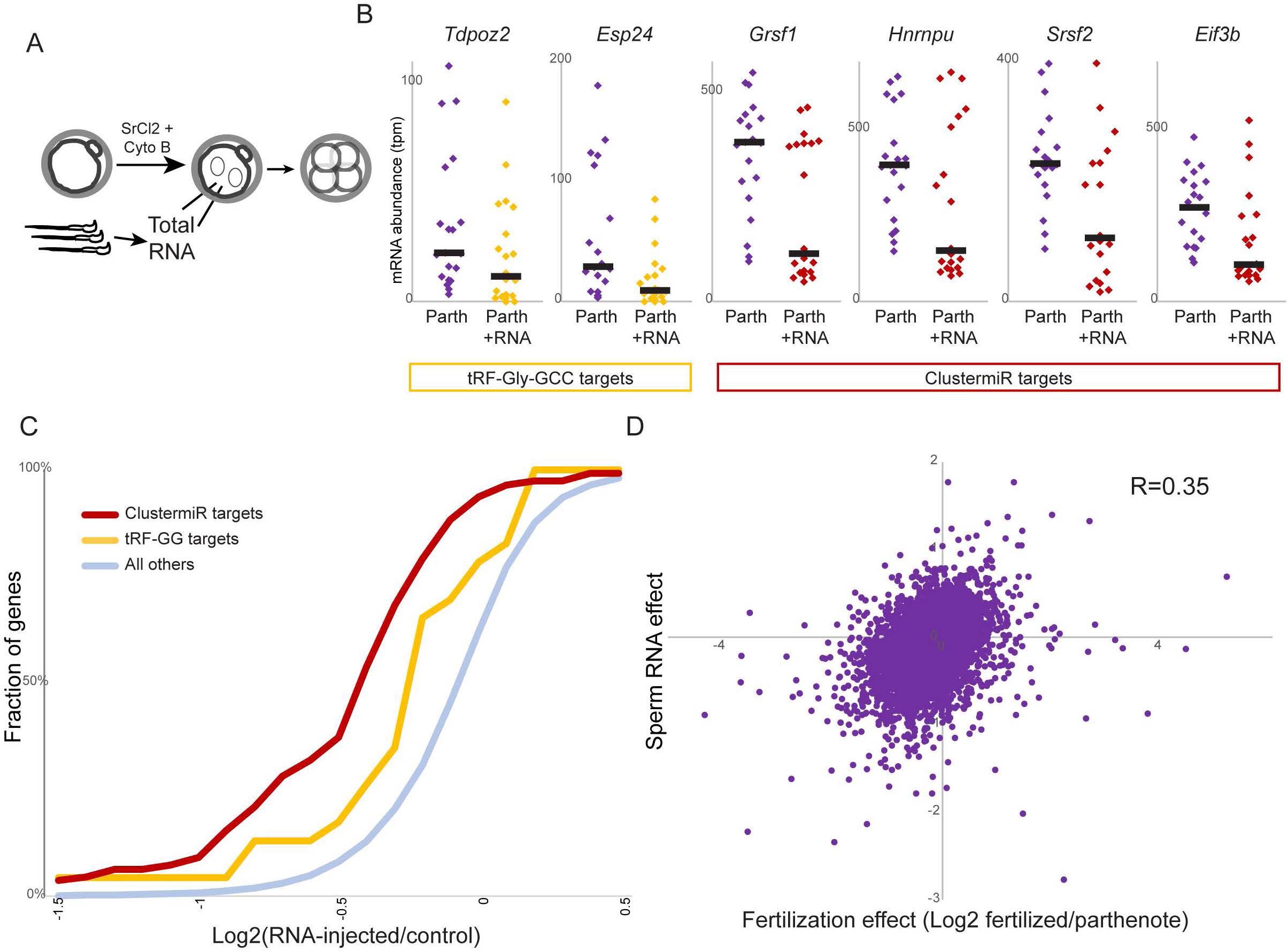
Parthenotes as a model system to investigate sperm RNA functions. A) Schematic of sperm total RNA injections into parthenotes. B) Dot plots showing sperm RNA-mediated repression of individual tRF-Gly-GCC target genes (Sharma et al., 2016) or clustermiR target genes (Conine et al., 2018, 2019) at the 4-cell stage. Note that none of these individual changes in mRNA abundance are statistically significant due to the inter-embryo variability in the RNA-injected parthenotes, which presumably reflects failed or inefficient injections. Factors leading to poor injections could include RNA degradation or clogged microinjection needles. C) Cumulative distribution graph showing change in expression in RNA-injected parthenotes (x axis) for tRF-GG targets or clustermiR targets, or for all remaining genes in the genome. Analysis here only includes genes expressed at a level of at least 10 tpm in either the parthenote or the parthenote+RNA data set, and so does not include maternally-silenced genes (Figure 4A) which are not expressed in either case. D) Scatter plot comparing the effects of fertilization (log_2_FC fertilized embryos/parthenotes, x axis) with the effects of sperm RNA injections into parthenotes (log_2_(parthenotes + sperm RNA)/(parthenotes + control RNA), y axis). The strong correlation documented here (R = 0.35) demonstrates that many of the gene expression defects observed in parthenotes are likely a result of the absence of key regulatory sperm RNAs.

Taken together, our analyses of gene expression in parthenotes further elaborate on our prior studies comparing embryos generated with sperm populations that differ in RNA payload. Parthenogenesis can therefore serve as a model system in which sperm RNA functions can be assayed on the “tabula rasa” of parthenotes developing in the absence of any sperm contributions.

### Embryo-to-embryo variation in preimplantation gene regulation

Perhaps the most novel feature of the dataset presented here is the genome-wide analysis covering hundreds of individual embryos, which stands in contrast to more typical efforts in which tens to hundreds of embryos are pooled prior to molecular studies. Our approach specifically allowed us to investigate variability in mRNA abundance across hundreds of embryos at a given developmental stage. Analysis of cell-to-cell variation across single *cells*, often leveraging GFP fusion constructs or single molecule FISH for analysis of individual transcripts, has been used in many studies to uncover determinants of noise in gene expression (Raj and van Oudenaarden, 2009). Here however, our data provide information on single *embryos*, averaging the output of tens of diploid cells. Our dataset is thus better considered a window on organismal “epivariation” (Youngson and Whitelaw, 2008), which may result from a multitude of contributing factors including:

1. genomic differences (XX vs. XY embryos, rDNA copy number variation, etc.)
2. differences in cell type composition of multicellular embryos, as in trophectoderm to inner cell mass ratio in blastocysts
3. differences in embryo viability or growth dynamics
4. cryptic parental effects from uncontrolled variables (intergenerational effects of paternal or maternal social status, for example)
5. stochastic epivariation (variability in spreading of cytosine methylation domains surrounding repeat elements, sperm-to-sperm variability in small RNA contents, variability of oocyte contents, etc.)
6. gene expression noise resulting from stochastic promoter activity (although this should be at least partially averaged out in multicellular embryos)
7. technical artifacts (resulting from, for example, batch effects)

There is a strong relationship between expression level of a gene and the variability in its expression (Bar-Even et al., 2006) driven partly by simple counting statistics – molecules present at low copy numbers exhibit higher coefficients of variation (standard deviation/mean). We therefore sought to define genes with unusually “noisy” or “quiet” expression across individual embryos, compared to other genes with the same mean expression level. After accounting for technical variation driven by sequencing depth and gene length (**Methods**), we characterized the relationship between mRNA abundance and gene expression variability. As a proof of concept, we initially analyzed datasets where we included both XX and XY embryos in the analysis, confirming the expectation that *Xist* and Y chromosome genes would exhibit unusually high variability in expression across embryos (**Figure 6-figure supplement 1**). This predictable finding validates the ability of our approach to identify genes with high variability between individual embryos, and motivated us to search for additional sources of variability in the dataset beyond sexual dimorphism. We therefore repeated this analysis in all the datasets in this manuscript, as well as several ICSI datasets from our prior studies (Conine et al., 2018), separating all the datasets based on the sex of the embryo. This yielded fifteen datasets each for the 4-cell and morula-stage embryos: parthenotes, five male/female pairs of datasets (1 NM, 2 IVF, and 2 ICSI cohorts) from this study, and two published male/female pairs of ICSI datasets (cauda sperm, testicular sperm) (Conine et al., 2018).

**Figure 6.**
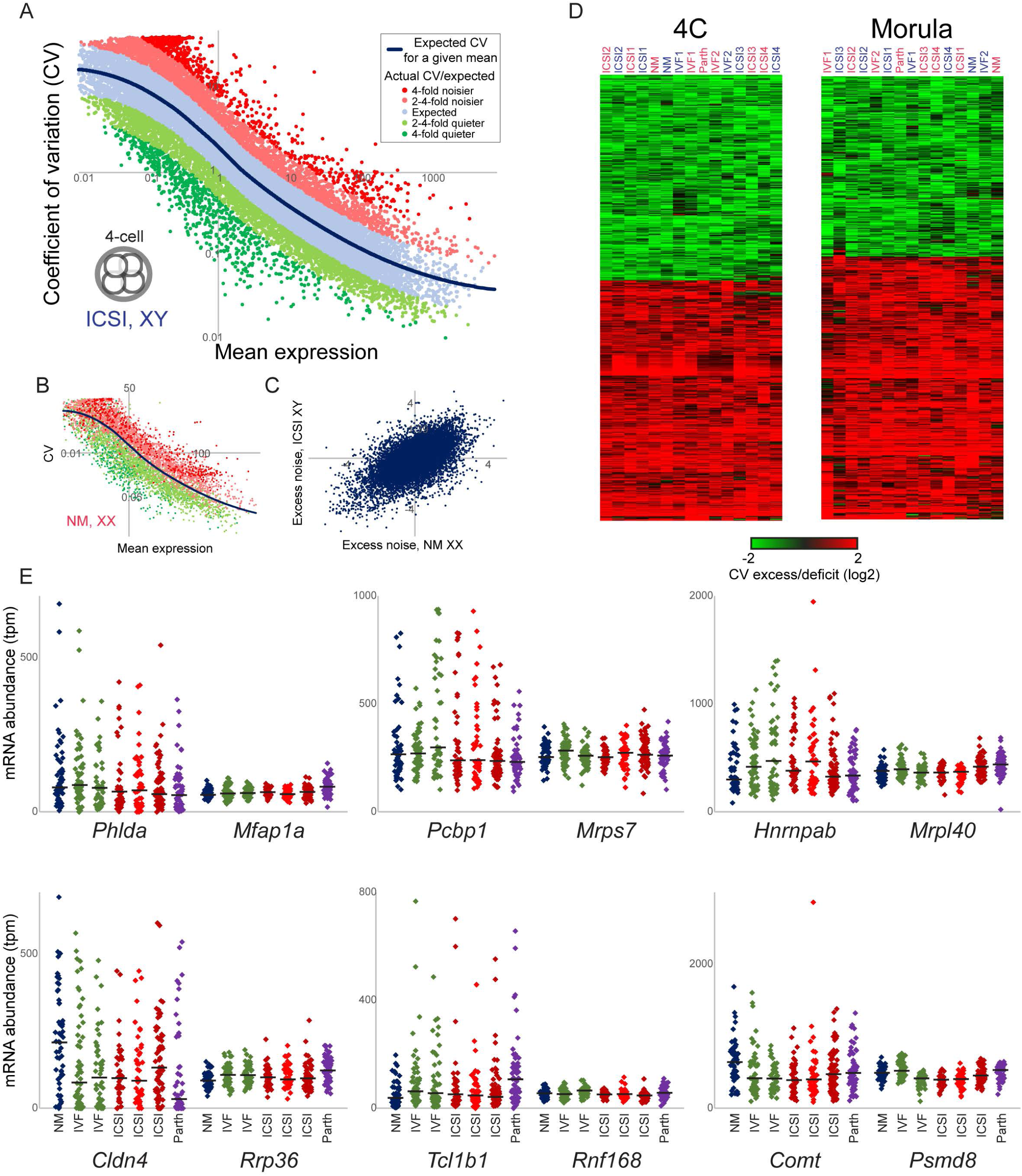
Embryo-to-embryo variability and constraint in gene expression. A) Global relationship between mean expression level (x axis) for all expressed genes, and variability in their mRNA abundance expressed as the coefficient of variation (CV, y axis), for a male ICSI four cell-stage data set. Blue line shows the global relationship between gene expression level and variability, and dots are colored according to their excess/deficit of variation (defined as log_2_ actual CV/expected CV). Importantly, although studies of gene expression noise differ in the measure of variability used – standard deviation, coefficient of variation (stdev/mean), and Fano factor (stdev^2^/mean) are all common – our goal is to identify genes exhibiting a relative excess or deficit of variation relative to other genes expressed at similar levels, and this is reasonably consistent across the three measures (not shown). We therefore used coefficient of variation for all analyses throughout, both because it is the most common measure of variance used in the literature, and because for most molecular readouts (eg, bimolecular interactions) relative changes in abundance are more relevant to function than absolute change in molecule numbers. B) Relationship between mean expression and CV for female natural mating four cell embryos, plotted as in panel (A). Here, only the noisy and quiet genes from the male ICSI dataset (A) are shown, colored as in that dataset, showing that the same genes are consistently relatively noisy and quiet across datasets. C) Scatterplot compares the noise “excess” (log_2_ actual CV/expected CV) for the male ICSI and female NM datasets from (A-B), showing all genes with a mean abundance >5 tpm. R=0.60. D) Measures of noise excess/deficit are highly reproducible across fifteen separate datasets for each developmental stage. Heatmap shows the residual variability (after correcting for an mRNA’s abundance) calculated using the indicated datasets, for all genes exhibiting at least a 2-fold excess or deficit of noise. E) Examples of high-variability genes along with lower-variance genes expressed at similar levels.

**Figure 6A** shows the characteristic relationship between mRNA abundance and gene expression variability for a typical data set – one of the male ICSI datasets – with the anticipated anticorrelation between mean expression and coefficient of variation (CV). Intriguingly, the overall relationship between RNA abundance and variation was nearly identical for ICSI, IVF, and NM embryos (compare solid blue lines in **Figures 6A-B**), despite the marked uncertainty (∼2-4 hours) in fertilization timing for IVF and NM embryos compared to the extraordinary temporal precision afforded by ICSI. This strongly suggests that little or none of the gene expression variation documented here results from differences in embryo timing.

To identify genes exhibiting unusually quiet or noisy expression relative to other genes expressed at similar abundance, we defined a “noise residual” as the ratio between the actual CV measured for a given gene, and the CV expected based on expression level (**Figures 6A-B**, blue line). **Figure 6A** shows genes colored according to this noise residual with red and green dots indicating relatively noisy and quiet genes, respectively. Remarkably, not only was the *overall* relationship between gene expression and variability consistent across datasets (blue lines in **Figures 6A-B**), but the noise residuals for *individual* genes were extraordinarily consistent across datasets. **Figures 6B-C** show a pairwise comparison between two datasets – one of the male ICSI datasets, and the female NM dataset. In **Figure 6B**, data from the female NM dataset are plotted for the noisy/quiet genes defined using the male ICSI dataset, showing that these genes are also noisy/quiet in the NM dataset. The scatterplot in **Figure 6C** more globally illustrates the remarkable reproducibility of the noise residual measure across these two datasets. This is extended to the entire dataset in **Figure 6D**, where noise residuals for all 15 datasets for each stage are clustered, with strong correlations between the residual noise identified in multiple independent datasets. In other words, by and large the same genes were stably-expressed, or epivariable, in morula-stage embryos whether generated by NM, ICSI, IVF, or parthenogenesis, and this variability was consistently observed in multiple distinct ICSI datasets collected over several years by several experimentalists. This is also readily appreciated by examining the underlying single-embryo data. **Figure 6E** shows individual embryo data for a number of unusually noisy genes (left panels), along with genes (ranging from typical variability to quiet) with roughly matching mRNA abundance, revealing consistent behavior across embryo cohorts (including parthenotes).

Along with other features of this dataset described below, these findings provide confidence that our single-embryo RNA-Seq data reproducibly quantifies biological variation in preimplantation gene expression.

### Features of constrained and variably-expressed genesets in the preimplantation embryo

We next sought to identify biological pathways enriched among the relatively quiet set of genes exhibiting low embryo-to-embryo variability. Unsurprisingly, but again supporting the validity of our approach, relatively constrained genes were highly enriched for those encoding basic housekeeping functions, with enriched gene ontology (GO) categories in both 4-cell and morula datasets including ribosome, translation, rRNA processing, and RNA splicing (**Figure 6-figure supplement 2**). These analyses are consistent with prior findings of constrained expression of housekeeping functions in single cells (Bar-Even et al., 2006; Newman et al., 2006), and extend this concept to the murine preimplantation embryo. In contrast, we find relatively few GO annotations, largely related to growth control, enriched among the most epivariable genes in our data set (**Figure 6-figure supplement 2A**). However, closer inspection (below) revealed several groups of biologically-coherent genes, each group associated with distinct GO annotations which are diluted when examining all epivariable genes jointly.

Are the unusually noisy genes markers of some underlying biological state of the embryo (such as the coherent differences in expression of scores of genes between XX and XY embryos – **Figure 1-figure supplement 1**), or do they exhibit uncorrelated noise across many individual genes which might result from promoter bursting, undersequencing, or stochastic “dropouts” due to technical failure to capture/amplify certain transcripts, etc? To address this question, we collated all highly-epivariable genes at each developmental stage, then for each embryo in the initial data set, we calculated each gene’s expression relative to all other embryos of that type (e.g. ICSI, IVF, etc.). Visualizing the correlation matrix between all epivariable genes (**Figure 7, left panels**) revealed substantial correlation structure at both developmental stages, with ∼3-6 groups of strongly co-regulated genes at each developmental stage. The relative expression of these genes across the individual embryos in our initial NM/ICSI/IVF/Parth cohort can be appreciated in the right panels of **Figure 7**. The co-regulation of large groups (up to ∼200) of genes is strong evidence that these variably-expressed genes are not simply technical artifacts resulting from stochastic dropouts or undersequencing, and instead argues that these genesets reflect embryo-to-embryo differences in some underlying state which manifests as coherent expression of regulatory modules.

**Figure 7.**
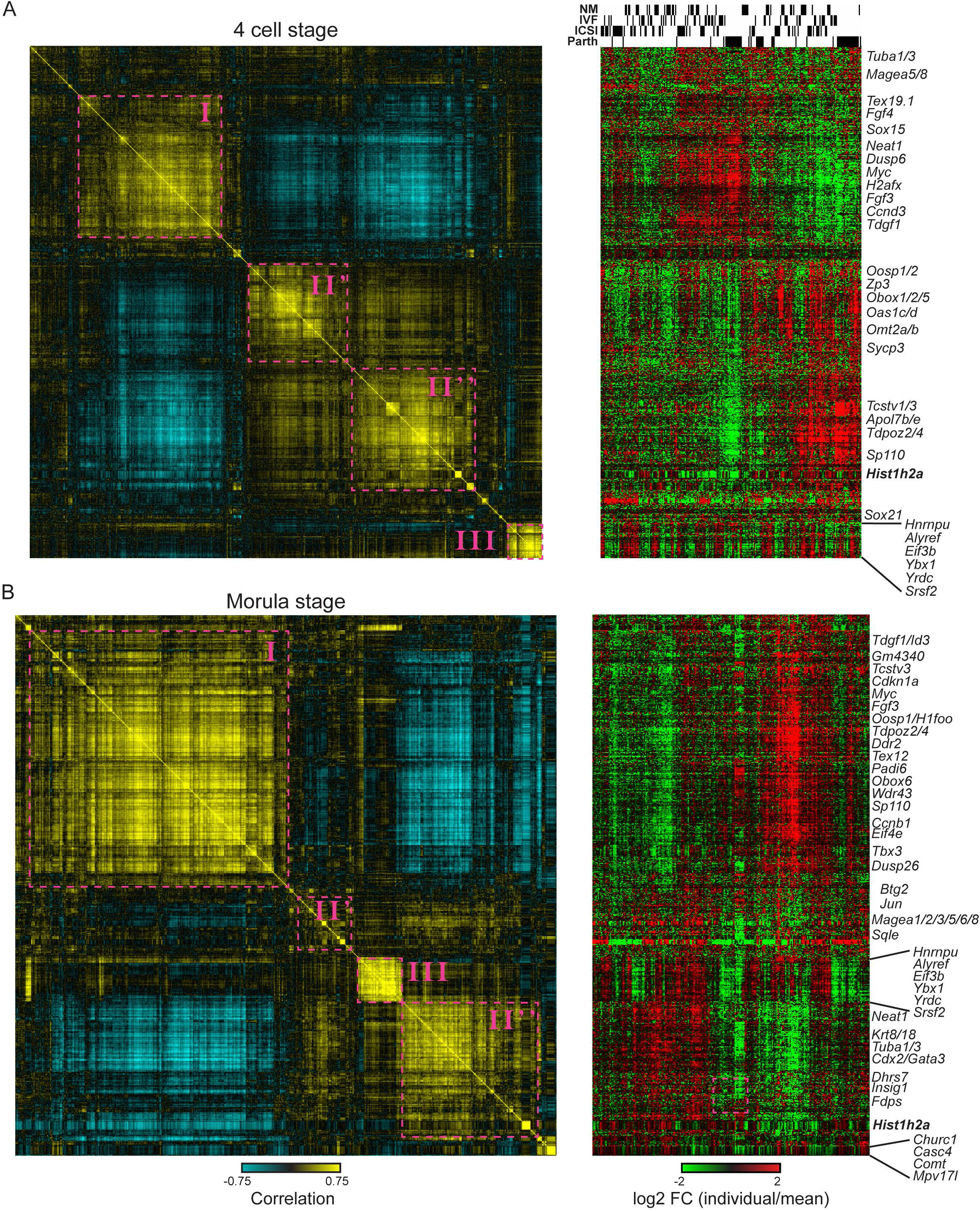
Co-regulation reveals coherent groups of epivariable genes in the early embryo. Epivariable genes for four-cell-stage (A) and morula-stage (B) embryos were defined as genes exhibiting an average two-fold excess of variance (CV) across all data sets (Figure 6D). For each stage, we show data here for the initial seven datasets (ICSI, IVF, and NM males and females, and parthenotes). For each gene, we calculated expression level compared to the median across the relevant dataset (e.g. ICSI embryos were compared to the ICSI median, etc.) as log_2_FC (sample tpm +1)/(dataset median tpm+1). Right panels show the resulting over- and under-expression in each embryo across our primary dataset, while left panels show the correlation matrix for all highly-epivariable genes examined here. For the 4-cell dataset, one cluster potentially reflecting cross-mapping was removed; the full dataset for all unusually noisy genes is shown in **Figure7-figure supplement 1**.

What is the biological significance of unusually-variable genes in the early embryo? A few interesting features stand out. At both developmental stages, clustering is dominated by two large anticorrelated clusters (I and II). In both cases, one of the clusters (I) includes Wnt-related signaling genes (*Fgf3*, *Tdgf1*, *Id3*, *Sox15*, *Tbx3*) and other genes involved in growth control (*Myc*, *Ccnd3*, *Cdkn1a*, *Prkcd*, *Tnip2*, *Mdm2*). At the four cell stage, these genes are anticorrelated in expression a large cluster (II) of genes which can be separated into subclusters enriched for oocyte-contributed transcripts (*Obox1/2/5*, *Omt2a/b*, *Oosp1/3*, *Sycp3*, *Oog3*, *H1foo*, *Gdf9*) and for MERVL-associated genes expressed upon zygotic genome activation (*Tdpoz2/4*, *Tcstv1/3*, *Sp110*, *Apol7b/e*). As both maternally-contributed transcripts, and ZGA-associated transcripts expressed during the late two cell stage, are being degraded during the 4-cell stage, it is plausible that Cluster II and Cluster I represent relatively early and late 4-cell embryos, respectively (although the high variability of these genes even in precisely-timed ICSI embryos makes this unlikely).

In morula-stage embryos, we again find a major separation of variably-expressed genes into two large groups (Clusters I and II) of anticorrelated genes (**Figure 7B**). Intriguingly, we again observe a large growth-related cluster (including *Myc*, *Fgf3*, etc.), which at this stage now also includes various maternal and ZGA-related transcripts. In other words, very early transcripts are anticorrelated with *Myc* expression in 4-cell embryos then become co-regulated with *Myc* in morulae. At this stage, expression of the Wnt/Myc-anchored cluster becomes anticorrelated with a cluster dominated by genes expressed in the trophectoderm lineage (*Cdx2*, *Gata3*, *Krt8/18*), tubulin genes (*Tuba1/2/3*, *Tubb2a/5/6*, *Tuba3a*, *Tuba4a*), and lipid metabolism genes (*Insig1*, *Pkm*, *Fdps*, *Sqle*, *Scd2*, *Fabp3*). Thus, similar to the behavior observed in four-cell embryos, morula-stage embryos exhibit two large populations defined by their expression of stage-restricted genes (here, the TE markers) in opposition to the Myc/WNT program. Whether this reflects developmental timing, or different cell cycle states (unlikely given the ∼16 unsynchronized nuclei present in the morula), or some deeper feature of early developmental growth control will be the subject of future studies. Importantly, although our data in principle would be explained by Cluster I transcripts being markers for the 8-cell embryo, examination of a full preimplantation gene regulation dataset (Conine et al., 2018) revealed that most of these transcripts are similarly-expressed in 4-cell, 8-cell, and morula-stage embryos, arguing against Cluster I/II distinctions representing a simplistic reflection of developmental timing.

Beyond the major division of embryos based on the Myc/Wnt modules, several smaller clusters are uncorrelated with the growth signature and appear to reflect independent underlying biological states of the embryo. Most remarkably, Cluster III at both stages is comprised of a large number of RNA and chromatin regulatory factors (*Hnrnpab*, *Hnrnpu*, *Alyref*, *Ybx1*, *Srsf2*, *Yrdc*, *Smarc5*, *Zmynd19*, etc.) whose expression is repressed in the embryo by the sperm-delivered clustermiRs (Conine et al., 2018, 2019). Smaller co-regulated clusters can also be appreciated, such as the co-regulated cluster (*Churc1*, *Casc4*, *Comt*, *Mpv17l*) highlighted in the morula dataset.

Taken together, our finding of high variability in the expression of a small number of regulatory modules identifies potential biological pathways subject to relaxed regulatory constraints in the murine preimplantation embryo.

## DISCUSSION

Altogether, this study presents a unique dataset comprising RNA-Seq data for 804 individual murine preimplantation embryos and parthenotes. We utilized this dataset to investigate three key topics: 1) the effects of assisted reproductive technologies on early embryonic gene regulation; 2) the use of chemically-activated parthenotes as a blank slate for interrogating sperm RNA functions in the embryo; 3) flexibility and constraint in early embryonic gene regulation. We note that this dataset has many other potential uses; for example, the RNA-Seq data cover entire transcripts and thus can be used to investigate splicing dynamics across early development.

### Persistent consequences of assisted reproduction methods

The era of assisted reproduction in humans began in 1978 and since then over five million live births have resulted from either IVF or ICSI. Thus, in addition to the subtle phenotypes documented in existing IVF cohorts (Halliday et al., 2014), it is important to note that potential effects of IVF or ICSI on complex diseases of later adulthood (diabetes, etc.) have yet to be assessed in human populations. These considerations have led to a substantial research effort using animal models to study potential long-term consequences of assisted reproductive methods (Doherty et al., 2000; Mann et al., 2004; Rinaudo and Schultz, 2004; Rinaudo and Wang, 2012; Rivera et al., 2008; Vrooman and Bartolomei, 2017). These studies have documented a wide array of metabolic and molecular sequelae of assisted reproduction that persist into adulthood, with glucose control and other metabolic functions being particularly susceptible to various perturbations ranging from embryo culture to the method of fertilization.

Here, we characterized effects of fertilization method on gene regulation in the preimplantation embryo, building on previous efforts from Rinaudo and others (Giritharan et al., 2012; Giritharan et al., 2010; Giritharan et al., 2007; Rinaudo and Schultz, 2004). Our study differs in several key details from prior efforts. First, we focused on developmental stages not characterized in most previous efforts, i.e. 4-cell and morula stages. Second, all of our embryos, including the natural mating cohort, were cultured in vitro under what are considered ideal conditions – KSOM growth media, 5% O_2_ tension. This protocol contrasts with prior studies that have typically compared in vitro cultured embryos to natural mating embryos which develop in the oviduct up until collection. Our study design compares embryos under identical conditions and thereby internally controls for the well-characterized contribution of embryo culture to gene regulation.

Overall, our findings are reasonably consistent with prior studies on the effects of fertilization on early developmental gene regulation. We found evidence that both IVF and ICSI-derived embryos exhibit altered expression of genes involved in mitochondrial function relative to NM embryos, consistent with prior studies (Giritharan et al., 2010). That said, in contrast, prior work from the same group found no significant changes in expression of nuclear-encoded mitochondrial genes in IVF blastocysts relative to NM blastocysts (Giritharan et al., 2007), although this is potentially explained by the fact that embryos in the 2007 study were cultured in suboptimal conditions (WM media at 20% O_2_, rather than KSOM at 5% O_2_). We also find that NM embryos express higher levels of ribosomal protein genes than either IVF or ICSI embryos, even at the morula stage, suggesting that some aspect of natural fertilization supports optimal growth of resulting embryos that persists for multiple cleavages following embryo collection and culture in vitro. Narrowing down the underlying basis for this altered gene expression program – whether it reflects cumulus cell or oviduct effects on the zygote, or differences between ejaculated sperm and sperm purified following epididymal dissection – may suggest modifications to assisted reproduction to more accurately model natural conception.

More interestingly, we found that the more extensive perturbations associated with ICSI compared to IVF are accompanied by greater alterations in embryo gene expression. Under our conditions, we found highly reproducible downregulation of trophectoderm markers in morula and blastocyst-stage embryos. Again, it will be valuable to uncover the inciting perturbation that drives these downstream molecular sequelae. Although our initial hypothesis was that piRNAs present in the sperm tail (Sharma et al., 2018) might play some role in this process, we found that injection of testicular piRNA populations had no effect on gene regulation in ICSI embryos.

Whatever the mechanical or molecular perturbation that results in this change in TE marker expression, we note that alterations in either TE cell number or gene expression are expected to affect placental development and ultimately drive metabolic changes in offspring. Identifying the reason for TE marker gene changes in ICSI embryos – and whether these changes are consistently observed across the various technically-similar but nonidentical ICSI protocols employed by different labs – will therefore be of great interest for optimizing human assisted reproduction.

### Parthenotes as a model for investigating sperm RNA functions in the embryo

Other than the haploid genome delivered by sperm upon fertilization, the zygote receives the vast majority of its molecular and epigenetic machinery from the oocyte. Yet it is increasingly clear that mammalian sperm can deliver functional RNAs to the zygote, as several studies have documented defects in preimplantation growth and development (Yuan et al., 2016) or in early embryonic gene regulation (Conine et al., 2018, 2019; Sharma et al., 2016) in embryos fertilized using sperm lacking (or supplemented with) specific small RNAs. Moreover, sperm RNAs have been implicated in mediating the effects of paternal environments on programming offspring metabolism and stress-related behaviors, as microinjection of various purified or synthetic RNAs has been shown to recapitulate a subset of paternal effect phenotypes in several studies (Chen et al., 2016; Gapp et al., 2014; Grandjean et al., 2015; Rodgers et al., 2015; Sarker et al., 2019).

To establish a model for analysis of sperm RNA functions, we therefore included parthenogenetically-activated embryos in this study. Our analyses are broadly consistent with prior molecular studies of uniparental murine embryos (Inoue et al., 2017a; Inoue et al., 2017b), readily identifying scores of genes that have been shown to be repressed by H3K27 methylation deposited in the oocyte. Notably, the methods used to generate uniparental embryos differ significantly between our study and Inoue *et al*, as Inoue *et al* generated fertilized zygotes and subsequently swapped paternal and maternal pronuclei to generate zygotes carrying two maternal (or paternal) pronuclei – creating gynogenetic (or androgenetic) embryos. As these embryos were generated from fertilized zygotes, they differ from the chemically-activated parthenotes analyzed here, which have not received any sperm contributions whatsoever.

Unsurprisingly, parthenotes exhibit massive changes in the early embryonic transcriptome, with the expected loss of expression of maternally-silenced genes being the most dramatic consequence of parthenogenesis. However, thousands of other non-imprinted genes are misregulated in parthenotes (**Figure 1B**), although many of these genes presumably reflect secondary changes resulting from imprinted gene misregulation – for example, overexpression of X-linked genes in parthenotes is a clear and predictable downstream consequence of decreased expression of *Xist* during the window of imprinted *Xist* expression (Inoue et al., 2017b).

Nonetheless, we show that a subset of gene expression changes in parthenotes result from the absence of sperm RNA delivery in this system. We have previously linked two distinct RNAs – a specific tRNA fragment, 5’ tRF-Glycine-GCC, and a group of clustered microRNAs present in testicular spermatozoa and cauda sperm but surprisingly absent from caput epididymal sperm – to specific gene expression programs in the preimplantation embryo (Conine et al., 2018, 2019; Sharma et al., 2016). Here, we confirm the clear prediction of these prior findings, which is that microinjection of total sperm RNA would drive repression of these two gene sets (**Figures 5B-C**). In addition to these important confirmations of prior predictions, we find a remarkable number of other genes whose misregulation in parthenotes is reversed by sperm RNAs (**Figure 5D**), suggesting that some 10% or more of gene expression changes exhibited by parthenotes result from the absence of regulatory RNAs delivered by sperm. This finding establishes chemically-activated parthenotes as a powerful model system for interrogating the functions of sperm RNAs in preimplantation gene regulation and development, and paves the way for future efforts to identify additional functional RNAs delivered by sperm to the zygote.

### Flexibility and constraint in preimplantation gene regulation

Finally, in contrast to the majority of prior genome-wide analyses based on pools of preimplantation embryos, our single-embryo dataset provides a unique perspective on embryo-to-embryo variation in gene regulation. Our analyses reveal that expression variability of individual genes was remarkably consistent across multiple independent datasets (**Figures 6D-E**), and confirmed the expected constraints on expression of major classes of housekeeping genes. Conversely, hundreds of genes exhibited consistently elevated variation relative to their expression level; in principle, the embryo-to-embryo variability documented here could result from:

1. Technical variation resulting from stochastic capture or amplification of specific transcripts (those with relatively short PolyA tails, for instance, or RNAs capable of forming extensive secondary structure) would lead to variability in expression across our dataset. Although counting noise resulting from undersequencing is another possible contributor to apparent gene expression variability, this is unlikely to explain the noise residuals here – on average, noise at low expression levels is controlled for when correcting for a gene’s mean expression level. Other common sources of technical variation include batch effects.
2. Biological variation driven at the single-gene level by factors such as slow promoter licensing, as observed in many single-cell studies of gene regulation.
3. Biological variation resulting from differences in precise embryo staging, as a gene undergoing rapid changes in expression level will appear highly variable if embryos from early and late in the gene induction/repression process are collected.
4. Epigenetic variability, whether stochastic (as in the case of *A^vy^* cytosine methylation domains) or environmentally-driven. In the latter case, unappreciated differences in the experiences of the oocyte and sperm donors could occur even under the highly controlled conditions provided in our animal facility; for example, multiple animals in the same cage could differ in social status.
5. Genetic variability, most clearly seen with the extensive sexual dimorphism in gene expression between XX and XY embryos (**Figure 1-figure supplement 1**), but also potentially encompassing common variation such as rDNA or other tandem repeat expansion/contraction (Gibbons et al., 2014).

Features of the highly-epivariable genes documented here strongly argue against the first three sources of variation described above. The co-regulation of many genes across our dataset (**Figure 7**) is inconsistent with noise resulting from stochastic capture/amplification of transcripts. Similarly, although promoter-specific bursting presumably drives some level of variation in gene expression in individual blastomeres, the co-regulation of large groups of genes, along with the fact that our single-embryo data set averages the output of multiple genomes (∼32, in the case of the morula-stage data set), both argue against promoter bursting being responsible for the epivariation observed here. Finally, imprecise embryo staging is unlikely to account for the findings reported here, both because our dataset includes precisely-timed ICSI samples, and because many genes exhibit a significant excess of noise at two stages of preimplantation development.

Thus, in our view, our dataset most likely reveals relatively stable differences in embryo state that result in coherent co-regulation of large sets of target genes. Two key questions arise from this hypothesis. The first is the molecular underpinning of embryo-to-embryo variation in a given dataset. For example, we find that the targets of sperm clustermiRs (Conine et al., 2018, 2019) are tightly co-regulated across control embryos, yet exhibit high embryo-to-embryo variability (Cluster III in both **Figures 7A** and **B**).

Epivariation in this gene set could, in principle, result from sperm-to-sperm variation in levels of the relevant clustermiRs, or could alternatively suggest that this biological pathway integrates multiple inputs including those present in the female germline. Supporting the latter hypothesis, we find that these genes are also significantly epivariable in parthenotes (**Figures 6D-E**), as well as in an unpublished oocyte dataset (not shown).

The second topic of interest regarding this epivariation is the question of whether embryos that differ in expression of a given gene set exhibit any long-term sequelae later in development. In other words, do embryos expressing low levels of, say, the *Myc*/*Fgf3*/*Sox15* cluster, ultimately give rise to nonviable embryos, or to offspring with unusually efficient glucose control, etc.? Addressing such questions will require generating reporters to enable sorting of high and low expression embryos for prospective long-term studies of offspring phenotypes.

## PERSPECTIVE

Altogether, the data presented here represent an unprecedented starting point for the interrogation of embryo-to-embryo variation in preimplantation gene regulation. Future studies will address: 1) the potential for modifications to assisted reproduction to minimize potential effects on offspring; 2) the identity of sperm-delivered RNAs that play functional roles in the embryo; and 3) the causes and consequences of variability in biological state revealed by examination of early embryonic gene expression across hundreds of individual embryos.

## ACKNOWLEDGEMENTS

We thank Nick Rhind, Marisa Bartolomei, Jaime Rivera, John Yeh, and Shaila Chauhan, as well as members of the Rando lab, for critical reading of the manuscript and insightful discussions. We thank Ana Boskovic for helping us optimize parthenote microinjections. This work was supported by NIH grant R01HD080224. C.C. was supported a Helen Hay Whitney Foundation Postdoctoral Fellowship.

## MATERIALS AND METHODS

### Animal husbandry

Six-week-old FVB/NJ mice were purchased directly from Jackson Laboratory. 24 hours after arrival at the University of Massachusetts Medical School LRB animal facility, animals were switched from chow to control diet AIN-93g (Bioserv). After being on control diet for two weeks, one male and one female were placed together to establish a breeding pair. The first three litters generated by a breeding pair were weaned at three weeks after birth and subsequently fed a control diet. These mice were used as donors for all natural matings as well as oocyte and sperm isolations.

All animal care and use procedures were in accordance with guidelines of the Institutional Animal Care and Use Committee.

### Superovulation

Female 7-12-week-old FVB/NJ mice were superovulated by intraperitoneal injection of 5 IU pregnant mare serum gonadotropin (PSMG) (ProSpec HOR-272) followed by 5 IU human chorionic gonadotropin (hCG) (Sigma CG10) 48 hours later.

### Embryo generation: Natural Mating

To generate NM embryos, females were superovulated as described above, and immediately after hCG injection the females were placed in a cage with a male mouse for breeding. Successful breeding was verified by checking for a mating plug ∼14 hours later. 18 hours after hCG injections bred females were euthanized and dissected, and zygotes were flushed into KSOM embryo culture media. To remove cumulus cells, zygotes were placed in KSOM containing 3 mg/ml hyaluronidase. After several minutes in hyaluronidase the eggs were washed 4-5 times in KSOM, and cultured in KSOM in a 37°C 5% O_2_, 5% CO_2_ incubator to develop. 16 hours after hCG injection was used as the time of fertilization; 4-cell embryos were collected 46 hours after fertilization, newly compacted morulae at 75 hours, and blastocysts at 96 hours (although blastocysts were always screened to be a similar size and stage of development – mid-blastocyst – before being collected).

To generate the initial cohort of matched-isogenic NM, IVF, ICSI embryos, males used for natural mating were removed from the bred female cage and housed alone for one week. Subsequently, males were euthanized and sperm was collected by removing the cauda epididymis, making an incision in the epididymal tissue and squeezing to release the contents. Sperm was then split into two groups: one prepared for IVF and the other for ICSI (below).

Siblings of females used for NM were superovulated (as above). Eggs were collected from the oviducts of the females 14 hours after hCG injection and separated into three groups in KSOM: IVF, ICSI, and parthenotes.

### Embryo generation: IVF

Females were superovulated as described above and then euthanized 13-14 hours after hCG injection. The female was then placed on a 37°C hot plate during ovary dissection. Oocytes were collected by perforating the ampulla using a needle and cultured in a drop of KSOM + 1mM reduced L-glutathione (Millipore MR-020P-5F, Sigma G6013) under mineral oil (Sigma M8410) at 37°C, 5% O_2_, 5% CO_2_. Sperm were collected by dissecting cauda epididymides from 8-15-week-old male mice. Gentle pressure was applied to epididymis tissue using forceps and sperm were allowed to swim out during a 45-minute incubation in HTF + 0.75mM methyl-β-cyclodextrin (Millipore MR-070-D, Sigma C4555) at 37°C. 1×10^6^ sperm were added to the KSOM drop containing oocytes, and gametes were co-incubated for four hours. After the co-incubation period, zygotes were washed in KSOM and cultured until collection at 46 hours (four-cell stage) or 75 hours (morula stage) after completion of IVF.

### Embryo generation: ICSI

For ICSI, sperm were separated into heads and tails as follows: sperm collected as above were spun at 10,000 x g, washed once with PBS, resuspended in 500 μl PBS, and then drawn through a 26G needle on a 1ml syringe between 20-30 times. The shearing force from being drawn in and out of the needle removed the sperm head from tail for the majority of sperm. The sperm were then washed twice in Nuclear Isolation Media (NIM) 1% Polyvinyl alcohol (PVA) and finally resuspended in 100-200 μl NIM 1% PVA for use in ICSI.

Oocytes removed from the ampulla as above were prepared for ICSI by placing them in KSOM containing 3 mg/ml hyaluranidase to digest the cumulus cells away from the oocytes (as above). After several minutes in hyaluranidase, oocytes were washed 4-5 times in KSOM and cultured in a 37°C incubator until injected.

To fertilize the oocytes by ICSI, dishes were made with separate drops of NIM with 1% PVA for washing the injection needle, and for sperm, and drops of FHM with 0.1% PVA for oocytes, and finally covered with mineral oil. ∼20 oocytes at a time were placed into the FHM 0.1% PVA drops for subsequent injection. Sperm heads were then picked and injected into the oocytes. After completion of the injections, the injected oocytes were maintained at room temperature for 5 minutes, washed four times in KSOM, and then cultured in KSOM in a 37°C incubator 5% O_2_, 5% CO_2_, for development. With ICSI the precise time of fertilization is known. 47 hours after fertilization, four-cell stage embryos were collected, 75 hours for morulae (visually screened for recent compaction), and 96 hours for blastocysts (also screened, for typical mid-blastocyst morphology).

### Generation of parthenotes

Oocytes were collected 15-18 hours after HCG injection, as described above for IVF. Cumulus cells were removed by placing the oocytes in KSOM + 1mg/mL hyaluronidase (Sigma H4272) for several minutes, then washing in KSOM. Oocytes were then transferred to a drop of KSOM + 10mM SrCl2 + 4mM EGTA + 5μg/mL cytochalasin B (Sigma 255521, E8145, C6762) under mineral oil and incubated for one hour at 37°C, 5% O_2_, 5% CO_2_. After the incubation, parthenotes were washed in KSOM and cultured until collection at 47 hours (4-cell stage) or 75 hours (morula stage).

### Sperm RNA Injections

Sperm were collected by dissecting cauda epididymides and vasa deferentia. The tissue was placed in a dish of Whitten’s medium and 3-5 cuts were made while applying gentle pressure with forceps to allow the sperm out. The tissue was then incubated at 37°C for 30 minutes to allow sperm to swim out. Sperm were collected and spun at 10,000×g for 5 minutes to remove debris. Sperm pellets were washed twice in PBS and incubated in 0.01% SDS, 0.005% Triton-X on ice for 10 minutes to lyse somatic cells. Pellets were then washed twice in PBS and flash-frozen

Total sperm RNA was extracted by adjusting the total volume to 120μl with water. 66.6 μl of lysis buffer (6.4M Guanidine HCl, 5% Tween-20, 5% Triton-X, 120mM EDTA, and 120mM Tris pH 8.0), 6.6 μl Proteinase K (Qiagen 19131), and 6.6 μl 1M DTT were added to the samples. The samples were then incubated, with shaking, at 60°C for 15 minutes on an Eppendorf thermomixer. One volume of water was then added and the sample was transferred to a phase-lock column. For phase separation, 400 μl of TRI Reagent (MRC inc) and 160 μl BCP (1-bromo-2 chloropropane, MRC inc) were added. The samples were then mixed by inversion 10-15 times, followed by centrifugation at 20,000×g for 4 minutes. Next, the aqueous phase was removed and mixed with 20 μg of Glycoblue (Ambion) and 1 volume of isopropanol, and samples were precipitated overnight at -20°C. Samples were spun down at 20,000×g for 15 minutes at 4°C and washed in 1mL cold 70% ethanol. Samples were allowed to air-dry and were reconstituted in nuclease-free water. To remove further contaminants from RNA, samples were treated with DNase I (Qiagen 79254) for 15 minutes at room temperature. A volume of 300μL nuclease-free water and 400μL phenol-chlorophororm-isoamyl (PCI) (25:24:1) was added (Invitrogen 15593031) to the samples in a phase-lock tube. Samples were spun at 12,000xg for 3-5 minutes at room temperature. The aqueous phase was used for a subsequent PCI extraction three more times. Finally, an equal volume of chloroform was mixed with the aqueous phase and spun down in a phase-lock tube 12,000xg for 3-5 minutes at room temperature. The RNA was precipitated in an equal volume of isopropanol for 2 hours at -80°C, washed in ethanol, and reconstituted in water as before.

One hour following the completion of parthenogenetic activation, parthenotes were transferred to a dish containing several drops of KSOM (Millipore MR-020P-5F) under oil, which was pre-equilibrated at 37°C, 5% O_2_, 5% CO_2_. Parthenotes were injected with either 100ng/μL H3.3-GFP mRNA or a mix of 37.5ng/μL H3.3-GFP mRNA and 62.5 ng/μL total sperm RNA, resulting in an injection roughly equal to the RNA content of one sperm. All injections were carried out using a Femtojet (Eppendorf) microinjector set to 100 hPa injection pressure, 7 hPa compensation pressure, 0.1s injection time, while the dish was on a warming plate set at 37°C. After injection, parthenotes were washed in KSOM and cultured until collection at 37°C, 5% O_2_, 5% CO_2_. Fluorescence of H3.3-GFP was used as a marker of injection at the two-cell stage.

### Single-embryo RNA-Seq

During collection single embryos were placed for lysis in 5 ul TCL buffer (Qiagen) containing 1% (v/v) β-mercaptoethanol and stored at -80°C until performing single embryo RNA-seq cloning using a Smart-Seq2 protocol (Ramskold et al., 2012). RNA was first isolated using RNA-SPRI beads (Beckman Coulter). cDNA was generated from full-length poly-adenylated mRNAs using Superscript II (Invitrogen) and PCR for 10 cycles to amplify the DNA. Adaptors for indexing samples and sequencing were added to cDNA fragmented by Tn5-mediated tagmentation using the Nextera XT kit (Illumina). Finally, pooled libraries (96 samples) were sequenced on a NextSeq 500 (75 cycles – high output).

### Data analysis

Single embryo RNA-Sequencing reads were mapped using an RSEM pipeline created using Dolphin – a sequencing pipeline design platform created by the University of Massachusetts Medical School Bioinformatics Core – which included removing microRNAs, snoRNA, rRNA, and PCR duplicates. Embryos with fewer than 10,000 detected transcripts were removed from the dataset. For most analyses we focused on mRNAs with a median abundance of at least 10 tpm across the dataset. DESeq was used for calling significant changes in mRNA abundance comparisons between groups.

### Identification of epivariable genes

We used estimated read counts based on RSEM output, then accounted for variation from Poisson sampling by taking into account library size of each sample. Our model assumes a mean/variance TPM for each gene and that the actual value in each one is multiplied by sequencing depth and gene length, and this serves as input to Poisson sampling. We use maximum likelikood to estimate the mean and variance for each gene. Predicted expression value per sample is the posterior based on the mean/var and actual observed counts. Mean and variance used for subsequent analyses are the mean and variance of these expected values.

For each dataset, the relationship between CV and mean mRNA abundance was calculated (blue lines in **Figures 6A-B**), and noise “excess” was defined as log_2_(actual CV/expected CV). Unusually quiet or noisy genes for further analysis were selected based on a noise excess/deficit of at least 2-fold, averaged across 15 datasets (shown in **Figure 6D**).

## SUPPLEMENTARY MATERIALS

### SUPPLEMENTAL FILE LEGENDS

**Supplemental File 1. Embryo RNA-Seq statistics.**

Each sheet shows sequencing statistics for the indicated dataset (annotated according to Supplemental File numbers 2-6). For each dataset, columns show total read number, rRNA- and Repeatmasker-mapping reads, unique genome-mapping reads, and duplicate read counts.

**Supplemental File 2. 4-cell-stage data set.**

Single-embryo RNA-Seq data for 238 embryos that passed quality control. Column headings include fertilization/embryo activation method (ICSI, IVF, NM, and Parth). Data are normalized to tpm (transcripts per million).

**Supplemental File 3. Morula-stage data set.**

As in **Supplemental File 2**, for 254 morula-stage embryos.

**Supplemental File 4. Replicate 4-cell-stage cohort.**

As in **Supplemental File 2**, for 133 embryos. Here, we collected three groups of embryos: IVF, ICSI (microinjected with GFP mRNA only), ICSI+piR (injected with GFP mRNA as well as gel-purified piRNAs from the testis).

**Supplemental File 5. Replicate morula-stage cohort.**

As in **Supplemental File 4**, for 139 morula-stage embryos.

**Supplemental File 6. Parthenote injection data set**

As in **Supplemental Files 2-5**, for 40 GFP-injected or sperm RNA-injected parthenotes cultured to the four-cell stage.

## SUPPLEMENTAL FIGURE LEGENDS

**Figure 1-figure supplement 1.**
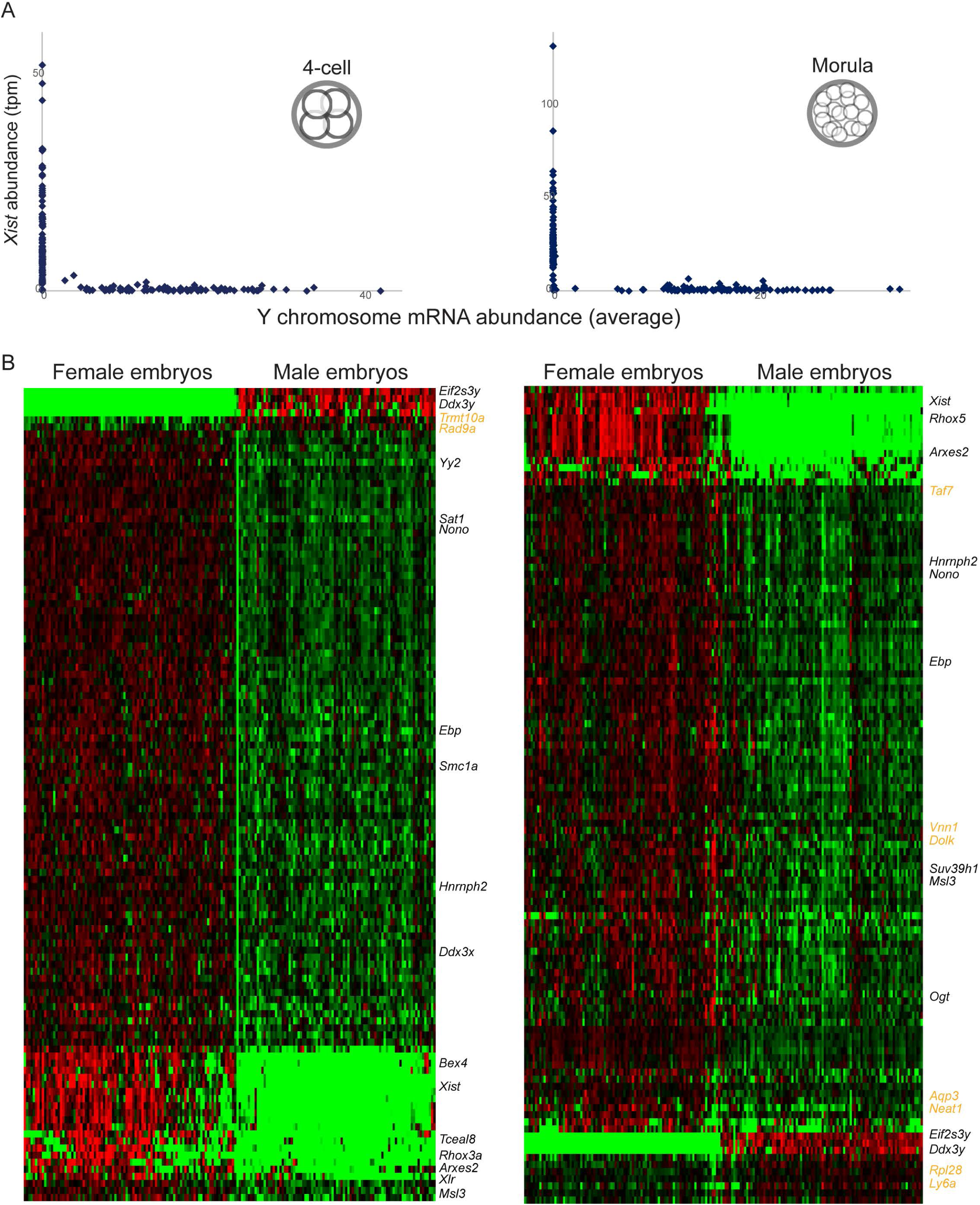
Separation of individual male and female embryos. A) Mutually-exclusive expression of *Xist* (y axis) vs. average expression of Y-linked genes (x axis) across all embryos in the 4-cell and morula-stage datasets. B) Sexually-dimorphic gene expression across the NM, ICSI, and IVF embryos. Heatmaps show genes exhibiting significant (multiple hypothesis-correct p value <0.05) differences between male and female embryos. Although the majority of these genes are X-linked genes exhibiting incomplete dosage compensation and thus female-biased expression, a small number of autosomal genes (names highlighted in orange text) also exhibit significant sexual dimorphism in our data set.

**Figure 2-figure supplement 1.**
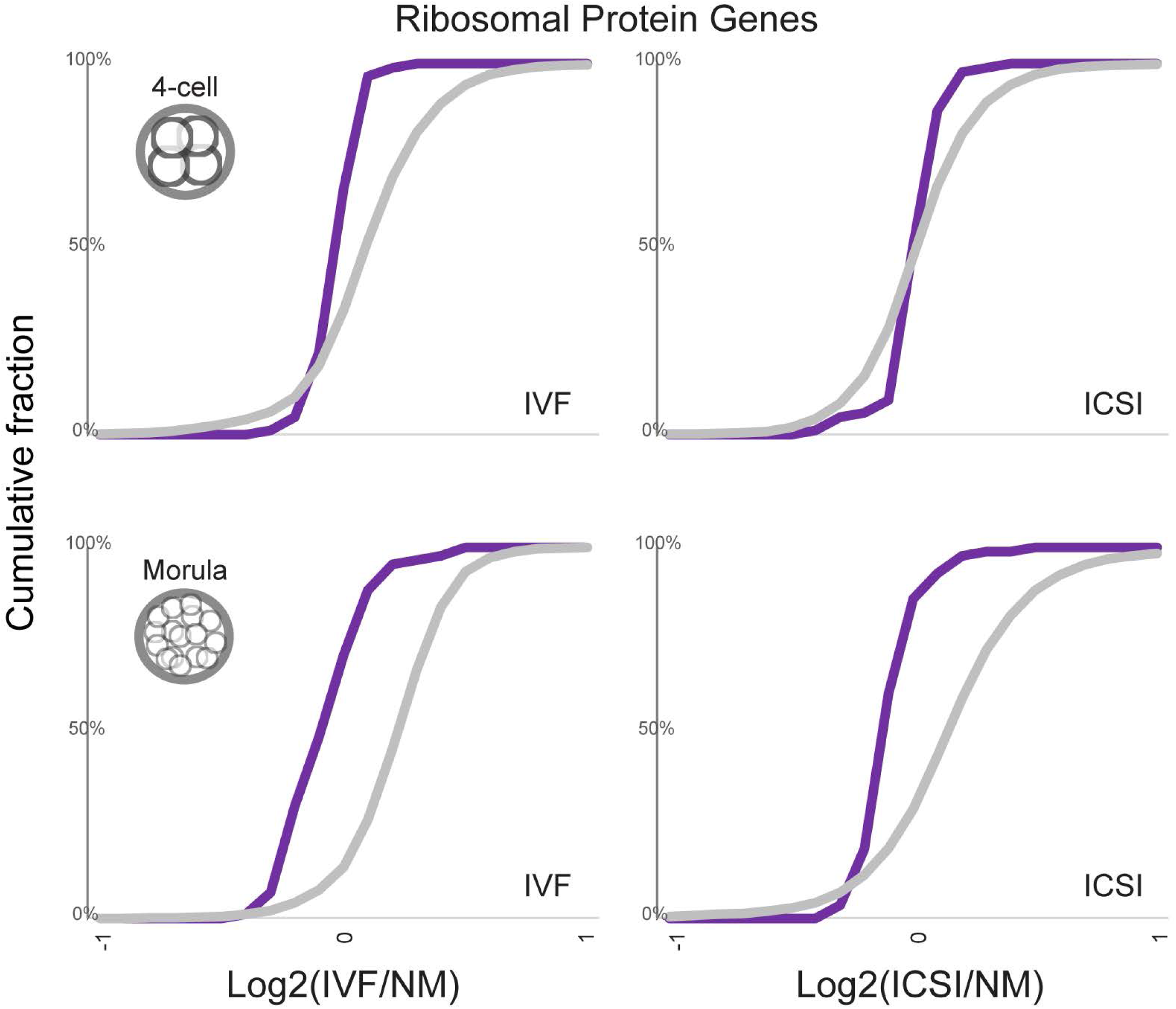
ART embryos exhibit downregulation of ribosomal protein genes. Cumulative distribution plots for ribosomal protein genes, and all other genes, showing the cumulative percent of genes (y axis) exhibiting the indicated expression changes (log_2_ (IVF/NM) or log_2_ (ICSI/NM), as noted on the x axis).

**Figure 3-figure supplement 1.**
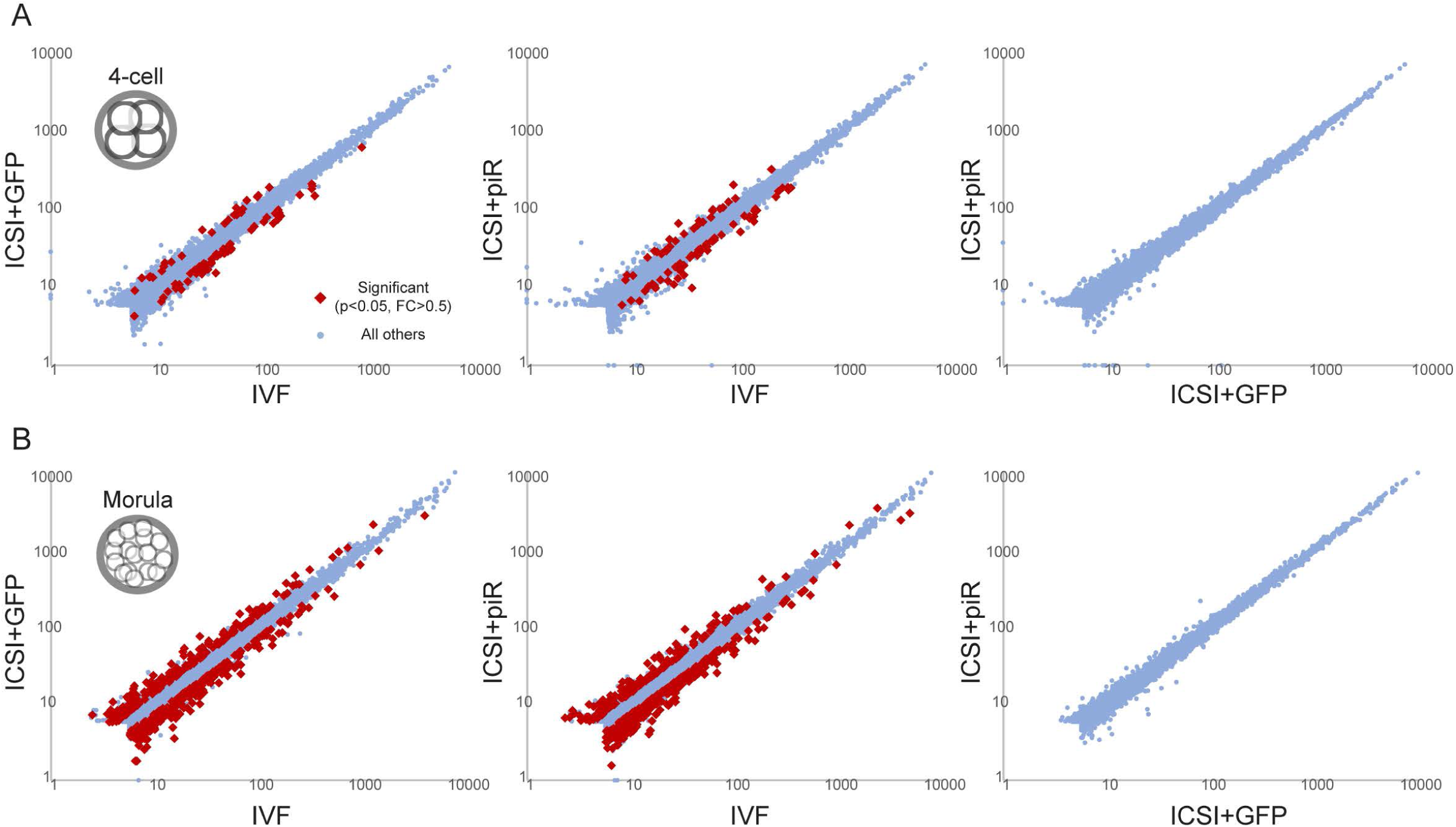
Collection of additional IVF and ICSI embryo cohorts. Scatter plots show pairwise comparisons between three new cohorts collected at the 4-cell (A) or morula stage (B). The three new cohorts include an additional IVF cohort and two ICSI cohorts subject to microinjection with either GFP mRNA alone (control) or GFP mRNA along with gel-purified (28-32 nt) testicular piRNAs. Interestingly, no genes were significantly affected by piRNA injection into ICSI embryos (see rightmost panels), essentially ruling out the possibility that failure to deliver piRNAs (due to the removal of sperm tails prior to ICSI) is responsible for gene misregulation in ICSI embryos.

**Figure 3-figure supplement 2.**
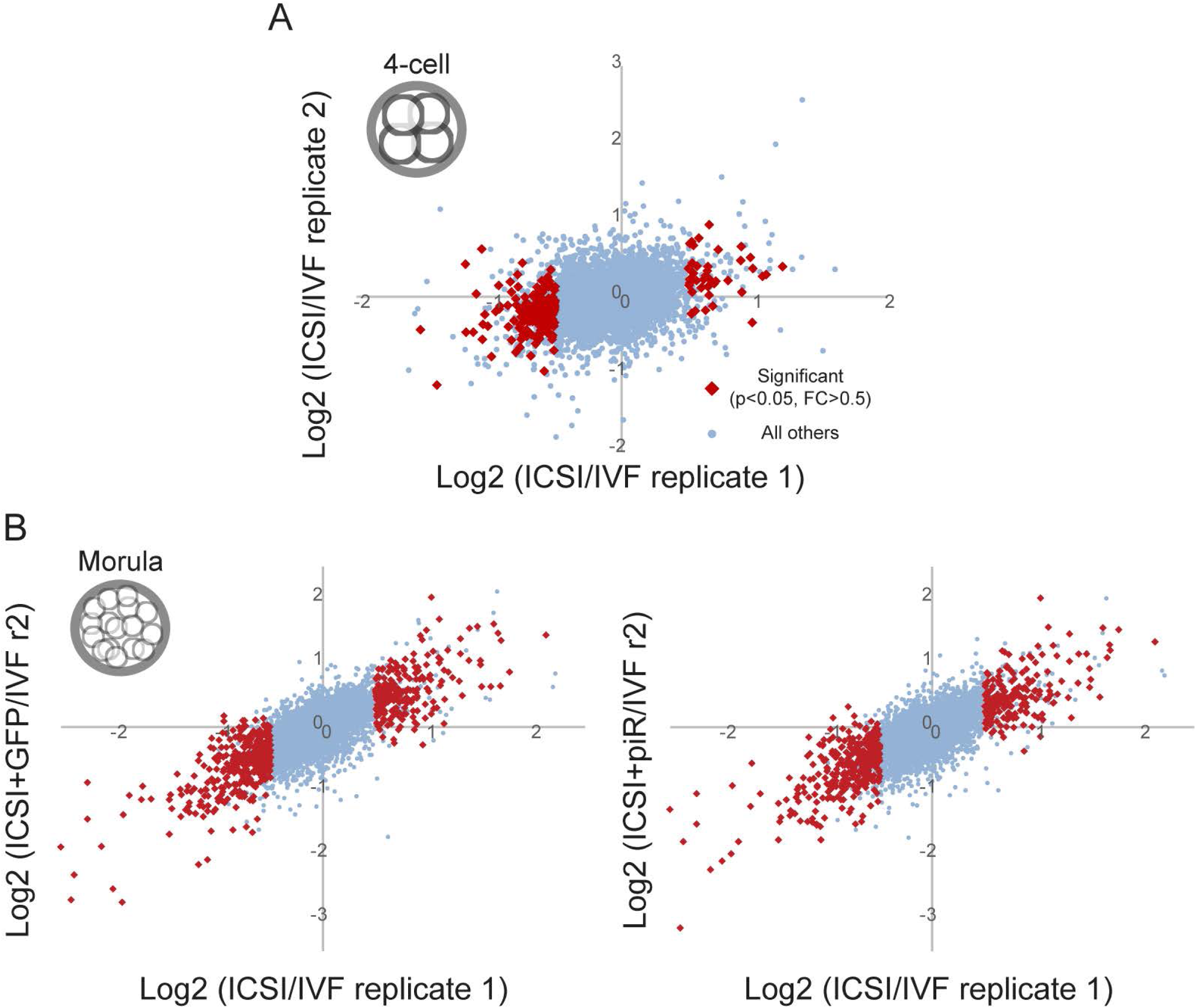
Reproducibility of ICSI effects on early gene expression. A) Four-cell-stage data. Scatter plot shows effects of ICSI relative to IVF, expressed as log_2_((ICSI+1)/(IVF+1)) for the original data set (replicate 1 – **Supplemental File 2**, x axis) compared to replicate 2 (**Supplemental File 4**, y axis). As the two new ICSI data sets (+GFP, +GFP+piRNAs) are indistinguishable, the ICSI value here is the median across the merged dataset. Red dots show genes that were significantly affected by ICSI in replicate 1. B) Morula-stage data. Scatter plots are shown as in panel (A), but here the two ICSI data sets are shown separately to further illustrate the reproducibility of ICSI effects on gene regulation.

**Figure 3-figure supplement 3.**
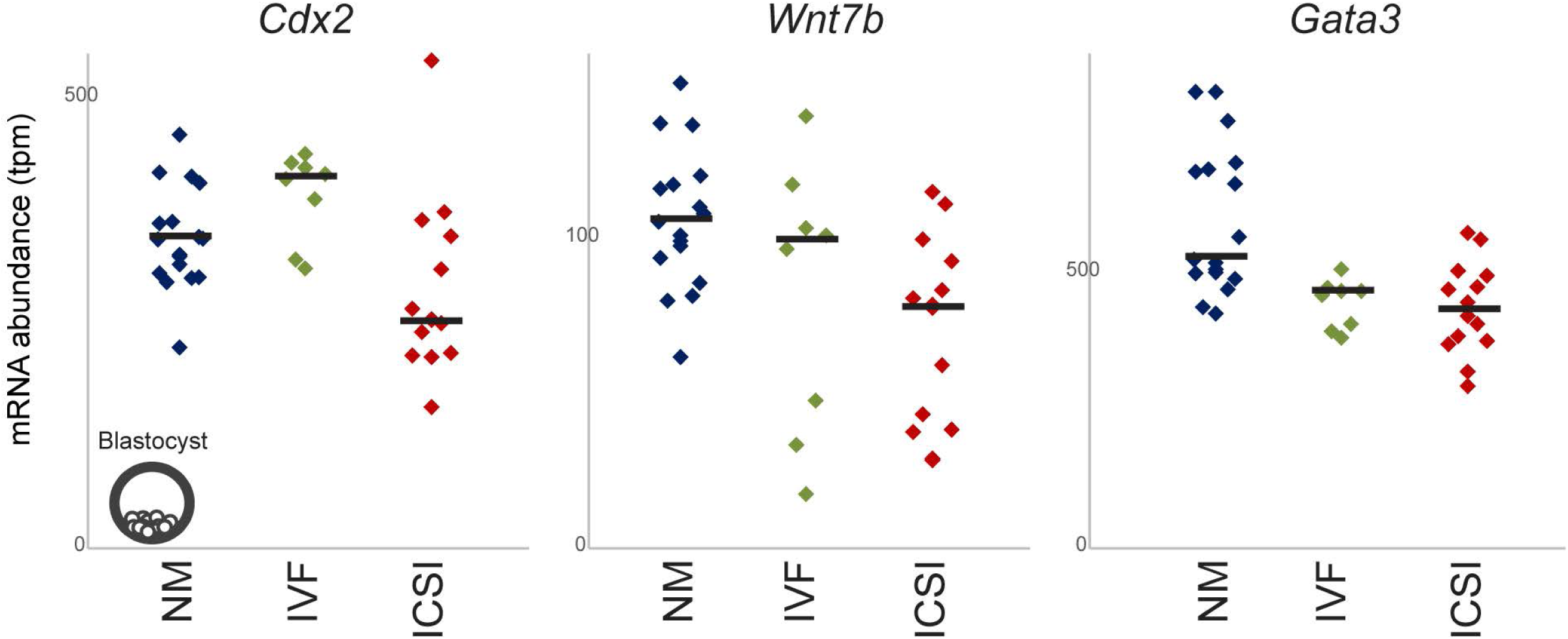
ICSI effects on TE marker gene regulation persist into the blastocyst. Dot plots for several trophectoderm markers in a small blastocyst-stage ART dataset.

**Figure 4-figure supplement 1.**
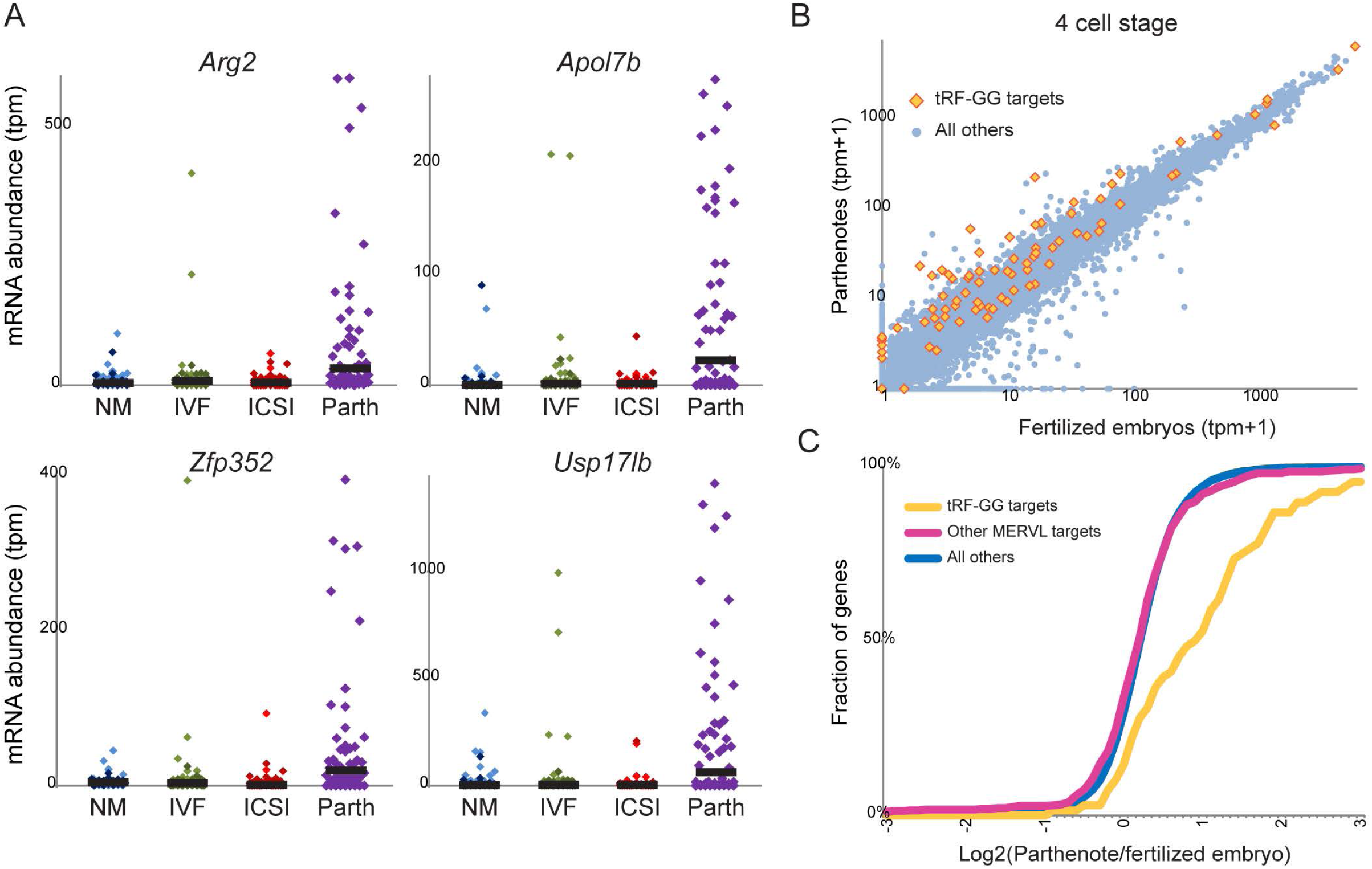
Overexpression of tRF-Gly-GCC targets in parthenotes. A) Dot plots for previously defined (Sharma et al., 2016) targets of tRF-GG in 4-cell-stage embryos/parthenotes. B) Scatter plot comparing parthenotes (y axis) with fertilized embryos (x axis), with tRF-GG targets indicated. C) Cumulative distribution graph showing change in expression in parthenotes compared to fertilized embryos (x axis) of tRF-GG targets, other MERVL-associated transcripts (Macfarlan et al., 2012), and all other genes.

**Figure 6-figure supplement 1.**
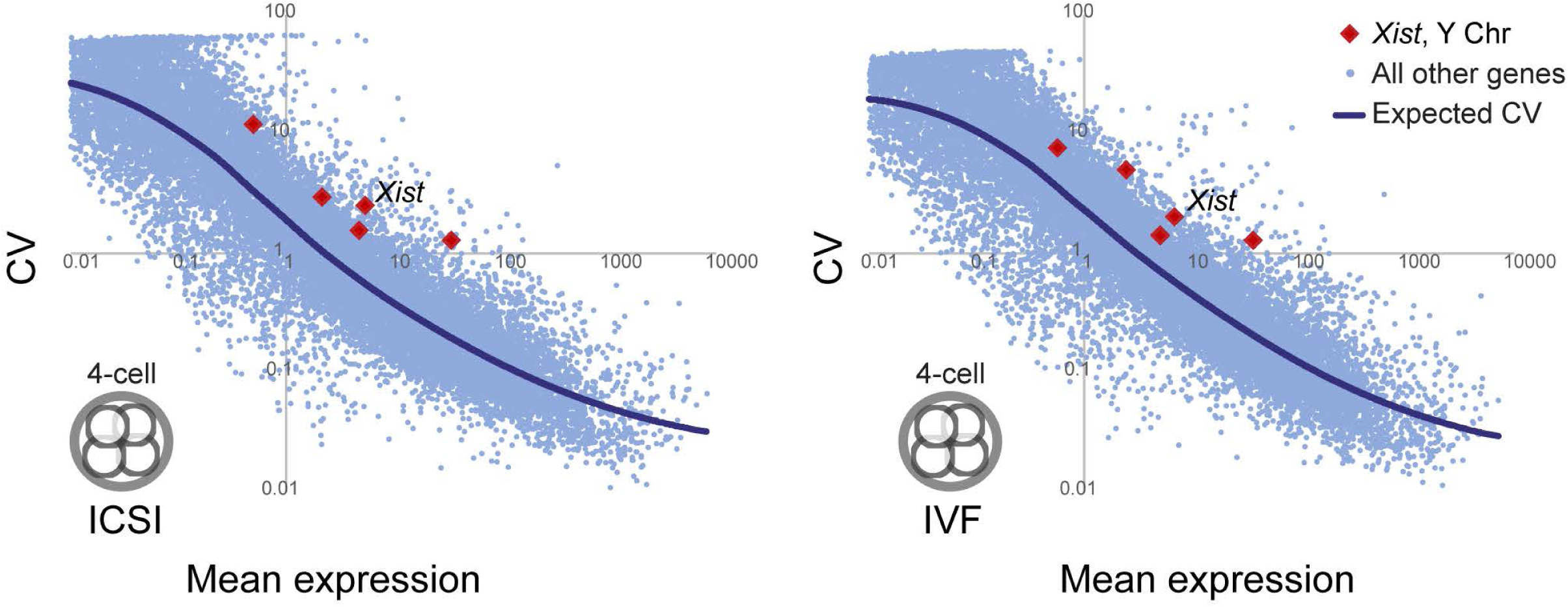
Gene expression noise across preimplantation embryos is reproducible. Analysis of variability across all ICSI embryos (left) or all IVF embryos (right) identifies high variation in sex-linked genes such as *Xist* (indicated). This confirms the power of this approach to identify biologically-relevant genes with high variability in a given data set, and motivates the separation of XX and XY embryos (see **Figure 1-figure supplement 1**) for the analyses of epivariation in Figures 6-7.

**Figure 6-figure supplement 2.**
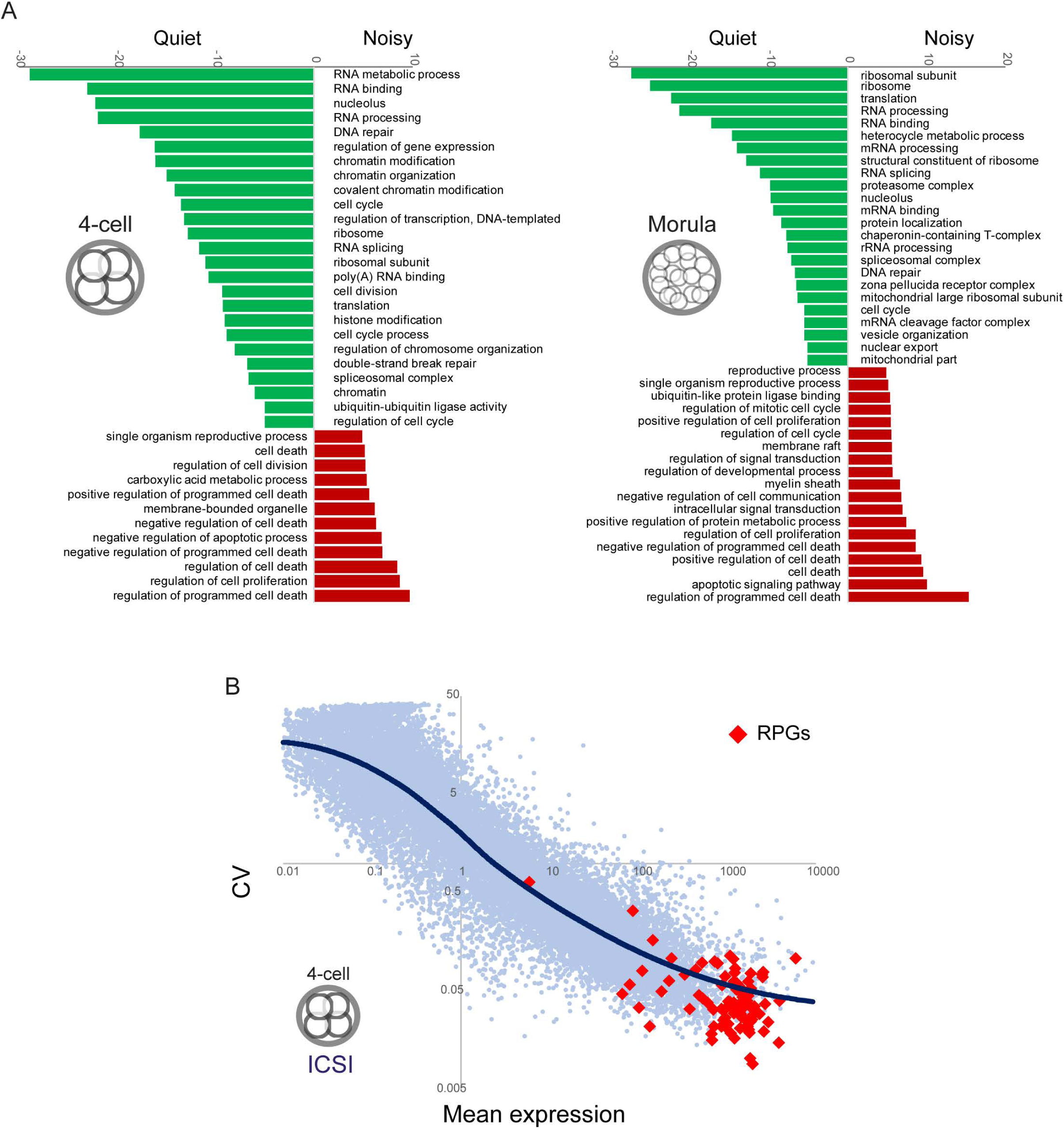
Biological features of quiet/noisy genes. A) Examples of GO categories enriched in relatively quiet or noisy genes, defined by genes exhibiting, on average across 15 independent datasets (Figure 6D), at least 2-fold lower or higher CV relative to that expected from mean expression. Bar plots show the -log_10_ p value for GO term enrichment, with enriched genesets for quiet and noisy genes, in 4C and morula embryos, as indicated. B) Ribosomal protein genes (RPGs) exhibit constrained expression in the early embryo. Relationship between Mean expression and CV for the four cell ICSI XY dataset is plotted as in Figure 6A, with RPGs indicated as red dots.

**Figure 7-figure supplement 1.**
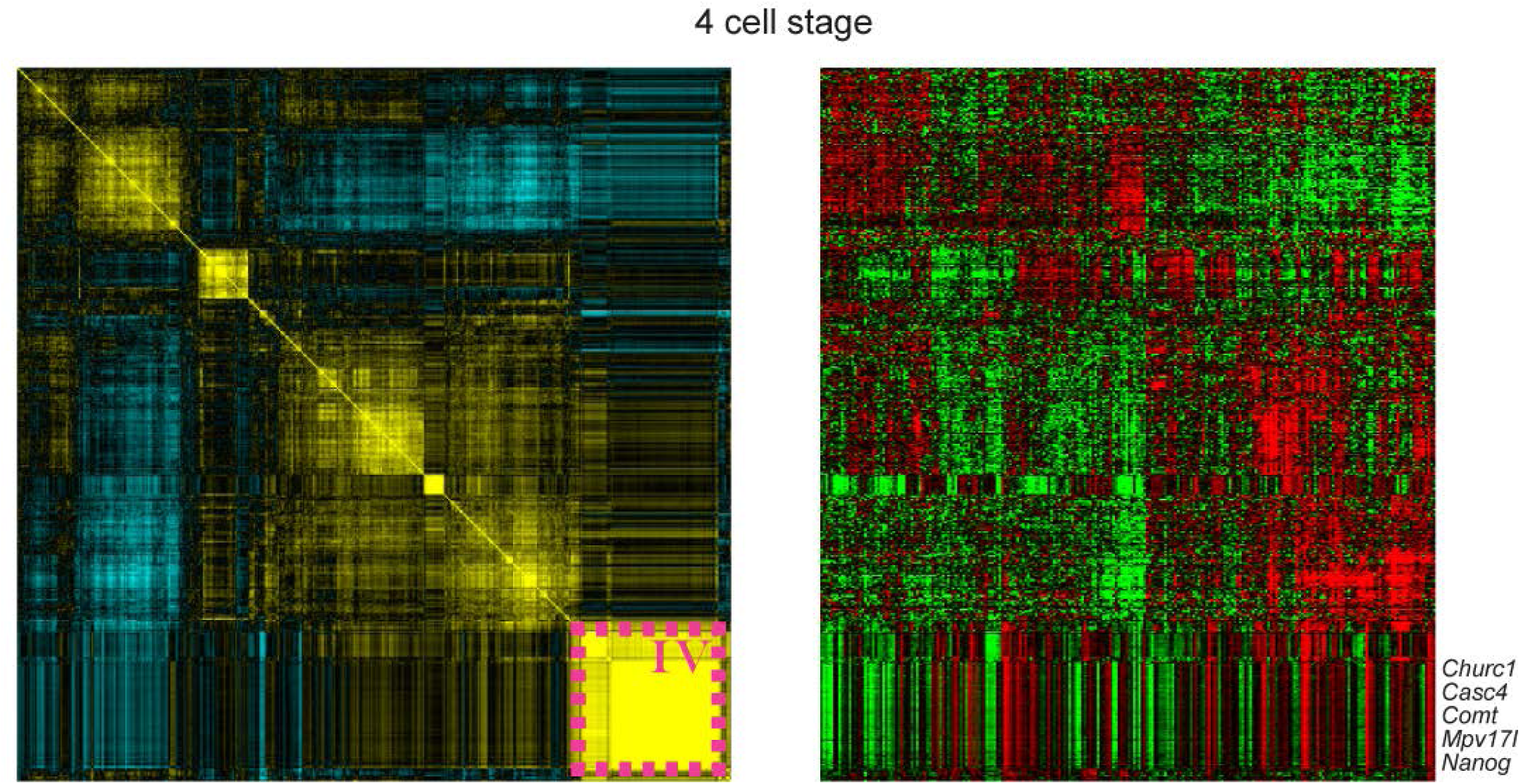
Full epivariation coregulation matrix for 4-cell embryos. Full correlation matrix and individual embryo gene expression values for all noisy genes at the 4-cell stage are reproduced here. Highlighted (“Cluster IV”) is a heterogeneous group of genes (not enriched for any GO annotations) that exhibit extraordinarily high correlations with one another, similar to those observed among clearly cross-mapping genes such as the gene cluster encoding Histone H2A. However, the genes in Cluster IV are not homologs, and visual inspection does not reveal any clear evidence for common repeat elements that would drive cross-mapping. We therefore removed these genes from the analysis in Figure 7A, but include them here.

